# Flexible perceptual encoding by discrete gamma events

**DOI:** 10.1101/2022.05.13.491832

**Authors:** Quentin Perrenoud, Antonio H. de O. Fonseca, Austin Airhart, James Bonanno, Rong Mao, Jessica A. Cardin

## Abstract

Spatiotemporal patterns of activity in the neocortex are linked to cognitive processes underlying behavior. However, identifying discrete underlying events within highly dynamic cortical network fluctuations remains a critical challenge. Here, we demonstrate a novel analytical method to track network events underlying state-dependent β- (15-30Hz) and γ- (30-80Hz) range activity in mouse primary visual cortex (V1). We find that γ events are selectively associated with enhanced visual encoding by V1 neurons and γ event rate increases prior to visually-cued behavior, accurately predicting single trial visual detection. This relationship between γ events and behavior is sensory modality-specific and rapidly modulated by changes in task objectives. These findings illuminate a distinct role for transient patterns of cortical activity, indicating that γ supports flexible encoding according to behavioral context.

---

Neural activity in the neocortex exhibits complex spatial and temporal patterns that dynamically reflect changes in behavioral state^1–6^. Moreover, disrupted activity patterns are a hallmark of many neurodevelopmental and psychiatric disorders^7,8^. Specific frequency bands of activity in the cortical local field potential (LFP), which arises largely from synaptic currents in local circuits, are linked to cognitive processes including attention, perception, and memory^9,10^. This patterned activity is often quantified as sustained oscillations^4,11–13^ but also commonly occurs in transient bouts that are difficult to detect reliably^14–19^. Establishing comprehensive links between spatiotemporal patterns of cortical activity and behavior thus requires novel approaches to detect and quantify discrete events, such as single cycles within a target frequency band, during dynamically regulated cortical activity.

To precisely identify discrete network events associated with state-dependent patterns of cortical activity, we recorded LFPs across cortical layers in primary visual cortex (V1) of freely running head-fixed mice (Fig. 1A-B, Fig. S1) and applied a novel analytical method for Clustering Band-limited Activity by State and Spectrotemporal feature (CBASS) (Fig. 1, Fig. S2, Supplementary Methods). CBASS first identifies candidate events in a single reference channel and sub-selects events whose laminar spectrotemporal profile is enriched in specific behavioral states (Fig. 1D, Fig. S2, Supplementary Methods).

**Figure 1.**
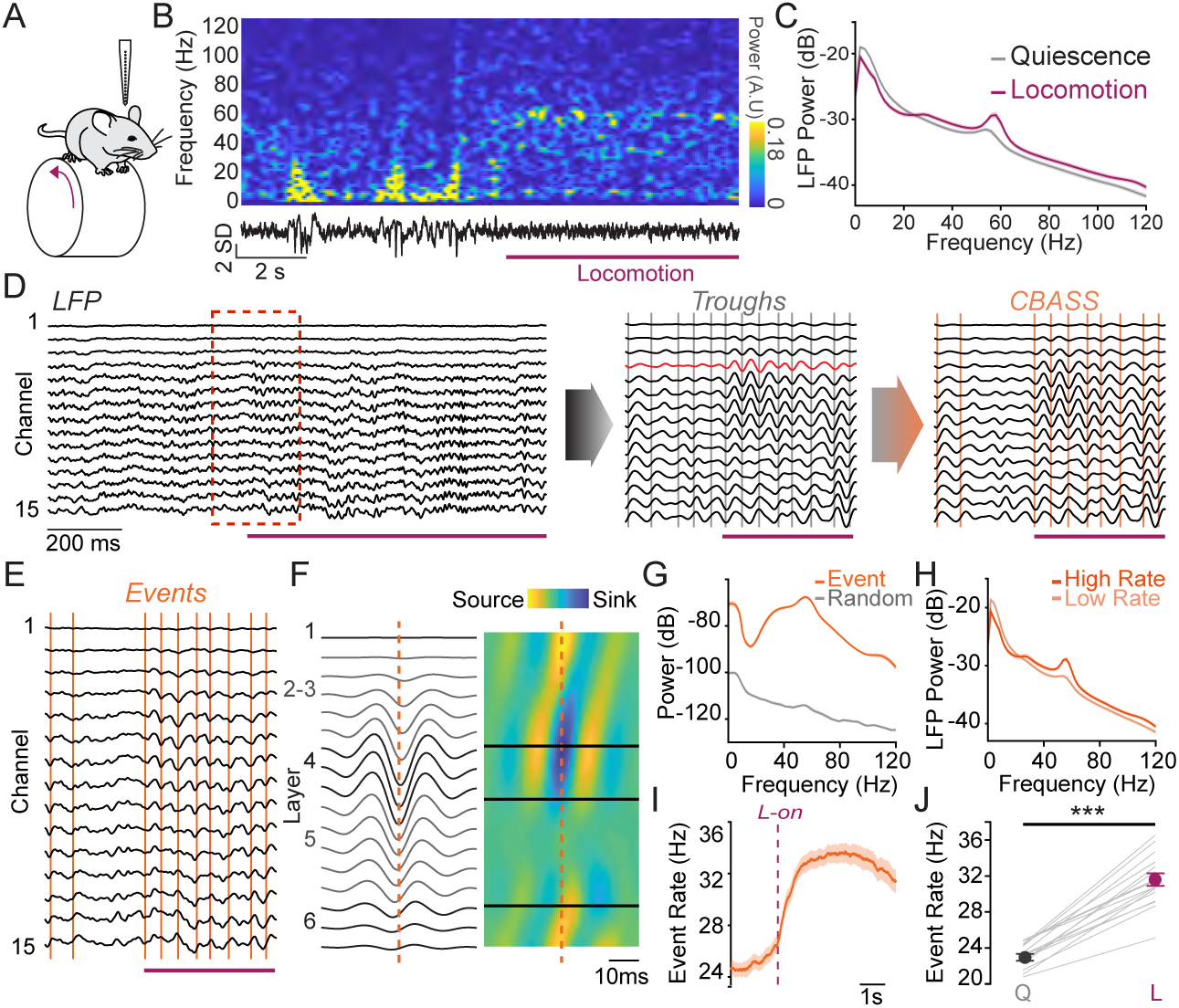
CBASS links global state-dependent changes in patterned activity to defined network events. **A**: Schematic of laminar cortical recordings in V1 of head-fixed mice on a running wheel. **B**: Example data showing one LFP channel and its short-time Fourier transform during a transition from quiescence to locomotion (purple). **C**: Average LFP power across channels (n = 19 mice), showing a selective power increase in the γ range (30-80Hz) during locomotion. **D**: CBASS applied to data from V1 during locomotion. *Left*: Multi-channel LFP. *Center*: Blowup of highlighted portion from left panel (red dotted lines), filtered in the γ (30-80Hz) range. Candidate events (gray bars) were selected at the troughs of the filtered signal in a reference channel (red). *Right:* Events (orange bars) whose spectral profile across channels are associated with that seen during locomotion are retained. **E**: LFP activity in the highlighted portion of panel D, showing events retained by CBASS. **F**: Average LFP around γ events (left) and associated CSD profile (right). Events are associated with a propagation of activity from layer 4 to superficial layers followed by deep layers. **G**: Power of the LFP events in F (orange) compared to matched random event averages (gray) (n = 19 mice). **H**: LFP power during high (upper quintile; dark orange) and low (lower four quintiles; light orange) γ event rate (19 mice). **I**: Rate of CBASS-detected γ events around locomotion onset (n = 19 mice). **J**: Event rate increases during locomotion (t-test; p = 8.26 x 10^−11^; n = 17 mice). Shaded areas: mean ± s.e.m. For detailed statistics see Supplemental Table 1.

A selective increase in γ (30-80Hz) power is observed in mouse V1 during locomotion^7–9^ (Fig. 1A-C), providing a well-defined context in which to examine discrete, repeated cortical network events in behaving animals. CBASS detected these γ events at a sustained rate in awake mice, suggesting that they are integral to awake cortical activity and coincide with propagation of activation from layer 4 to layers 2-3 and 5 (Fig. 1D-E)^20,21^. Detected events held considerable energy in the γ range (Fig. 1G) and had a stable current source density (CSD) profile (Fig. S3). LFP power increased in the γ range during high event incidence (Fig. 1H) and event rate increased 1.36 ± 0.1 fold during locomotion (n = 17 mice; Fig. 1I-J). All statistical results are listed in detail in Supplemental Table 1.

In addition to γ associated with locomotion, mouse V1 exhibits other prominent modes of patterned activity, including robust visually evoked β/low γ oscillations (hereafter referred to as β)^22,23^. γ and β arise from different excitatory-inhibitory interactions in the local cortical circuit^1,8,22,23^. In contrast to γ events, CBASS-detected events in the β range evoked by visual stimuli (Fig. S4A-C) had distinct profiles (Fig. S4D-F). β event rate increased 2.54 ±1.31 fold selectively during visual stimulation (n = 17 mice; Fig. S4G-H). β and γ events were interleaved on a fast time scale, indicating rapid switching of non-overlapping network processes (Fig. S5A-D). Co-labelling between γ and β events was limited (Fig. S5E), suggesting that CBASS resolves concurring categories of band-limited events in the cortex.

To examine the relationship between network events and subthreshold activity in individual neurons, we performed whole-cell patch-clamp recordings in cortical layers 2 to 5 of awake mice while simultaneously monitoring LFP across layers (Fig. 2A-B, Fig. S6). γ events coincided with rapid deflections of the membrane potential (Vm) riding on a slower overall depolarization^17,24,25^ (Fig. 2B-C, Fig. S7B, G, L). Events occurred with increased Vm power across frequencies (Fig. S7C, H, L) and a selective increase in Vm-LFP coherence in the γ band in all layers (Fig. 2D-E, Fig. S7D, I, N). γ events were precisely timed relative to spiking in all layers and were associated with a marked increase in spike-LFP synchrony in both intracellular (Fig. S7E, J, O-Q) and extracellular recordings (Fig. 2F-G, Fig. S8). γ event-associated spikes occurred earliest in L4 and latest in L2-3, consistent with feedforward thalamocortical processing (Fig. 2F). Synchrony was strongest in L2-3 (Fig. 2G, Fig. S7E) and markedly enhanced in fast-spiking (FS), putative inhibitory units relative to regular spiking (RS), putative excitatory units (Fig. S8D), in good agreement with previous reports^1,2,26^.

**Figure 2.**
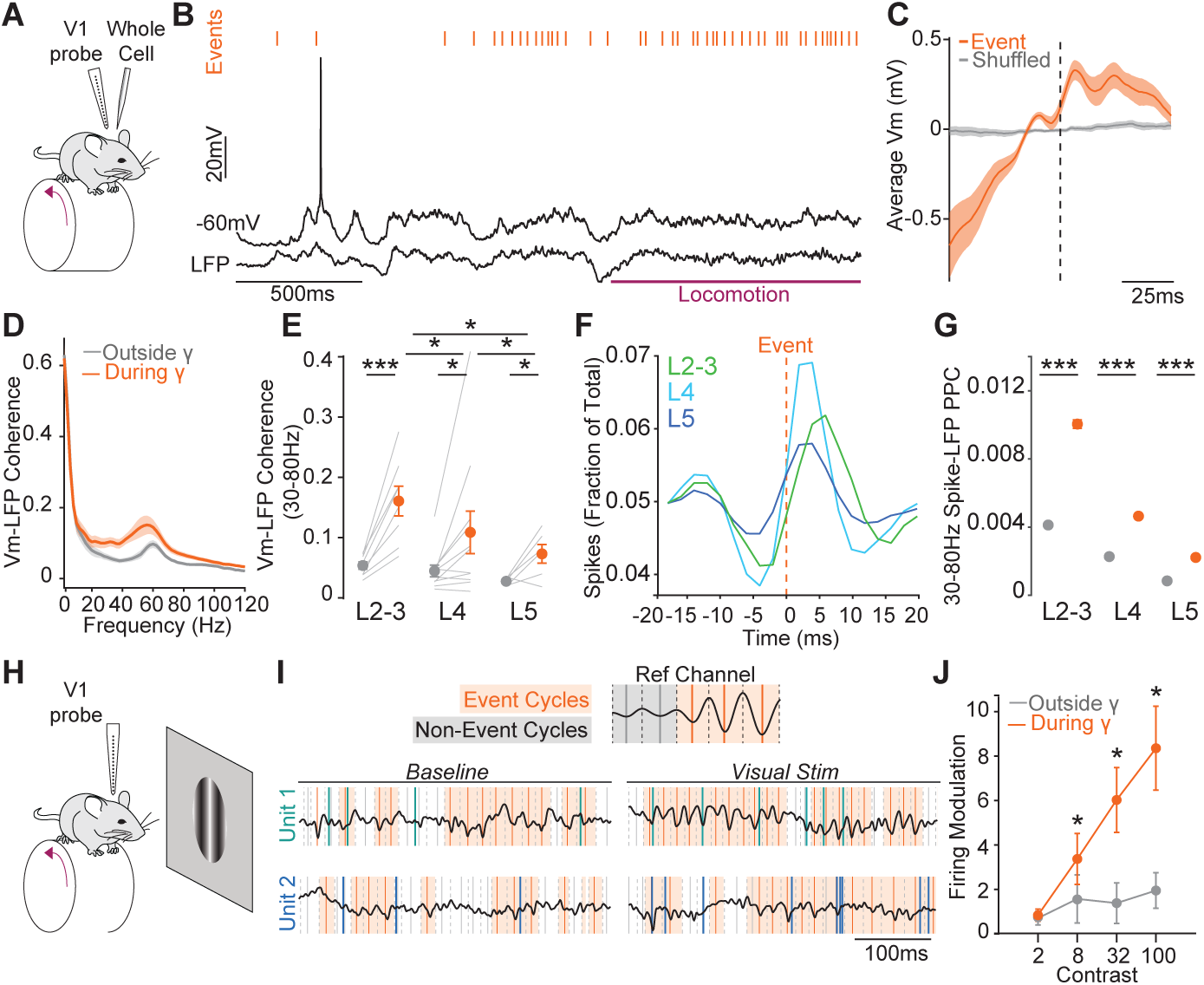
Network events regulate spike timing and enhance visual encoding. **A**: Schematic of simultaneous whole-cell patch clamp and laminar recordings. **B**: Membrane potential of a Layer 4 neuron, inverted LFP, and γ events (orange) around locomotion onset (purple). **C**: Average Vm around γ events (n = 25 neurons). **D**: Coherence spectra of Vm and LFP during (orange) and outside (gray) γ event cycles (n = 25 neurons). **E**: Overall γ coherence (30-80Hz) during (orange) and outside (gray) γ event cycles. (paired t-test within layers and unpaired t-test across layers). **F**: Population average distribution of spikes around γ events for neurons in layers 2-3 (green), 4 (cyan) and 5 (dark blue). **G**: Overall spike-LFP Pairwise Phase Consistency (PPC) in the 30-80Hz range, during (orange) and outside (gray) γ event cycles for RS units (Layer 2-3: 82 units; Layer 4: 68 units; Layer 5: 279 units; Welch’s t-test). **H**: Schematic of laminar recordings during retinotopically aligned visual stimulus presentation. **I**: Upper: Schematic of the analysis of spikes occurring during and outside γ events. Lower: Example activity of two V1 RS units before and after stimulus onset, illustrating visually evoked spikes during γ event cycles. **J**. Modulation of firing response to grating stimuli of increasing contrast during (orange) and outside (gray) γ event cycles (n = 47 RS units). Error bars: mean ± s.e.m. *α ≤ 0.05, *** α ≤ 0.001 For detailed statistics see Supplemental Table 1.

Spike-LFP synchrony within γ event cycles increased greatly during visual stimulation (Fig. S8). Event occurrence and RS unit spiking were uncorrelated during spontaneous activity but became correlated during presentation of high-contrast drifting gratings (Fig. S9), suggesting that visually evoked spikes occur preferentially during γ events. We therefore examined visual responses during and outside of γ events (Fig. 2H-I). We found that visual stimulation evoked almost no modulation of RS unit firing outside of γ event cycles (Fig. 2J, Fig. S10C). However, evoked firing was strongly enhanced during γ event cycles, regardless of behavioral state (Fig. 2J, Fig. S10C, E, G). Surprisingly, there was no similar enhancement during β events, despite their strong modulation by visual stimulation (Fig. S10A-B). Visually evoked spikes are thus selectively aggregated during γ events.

To examine the relationship of γ and β events to visually guided behavior, we trained mice in a visual contrast detection task (Fig. 3A, Supplementary Methods) that relies on V1 (Fig. S11) and shows behavioral state-dependent performance (Fig. S12). During hit, but not miss, trials γ event rate exhibited a consistent upward trajectory starting after stimulus onset (Fig. 3, Fig. S13) and peaking around lick response onset (Fig. 3B-D, Fig. S13C,E,G). In contrast, β event rate was unaffected by trial outcome (Fig. S13B,D,F,H). We performed a logistic regression to predict behavioral responses using γ and β event rates in specific time windows around the stimulus and response onsets (Fig. 3E). Prediction accuracy increased as the animal approached the lick response time. Deviance increase, parameter shuffling, and coefficient values indicated that γ, but not β, rate was critical for predicting trial-by-trial behavior (Fig 3G-H, Fig. S14A-J). These results were maintained when the analysis was restricted to periods of quiescence, indicating that they were not due to locomotion-associated γ events (Fig. S14K-O). Whisking did not modulate γ event rate, further indicating that the relationship between γ events and behavior is not simply the result of motor movements (Fig. S15). γ rate increases and model predictions were also significant during false alarm trials (Fig. S13E,G, Fig. S14F-J), suggesting that increased γ event rates anticipate task-relevant behavioral responses independent of visual stimulation or reward.

**Figure 3.**
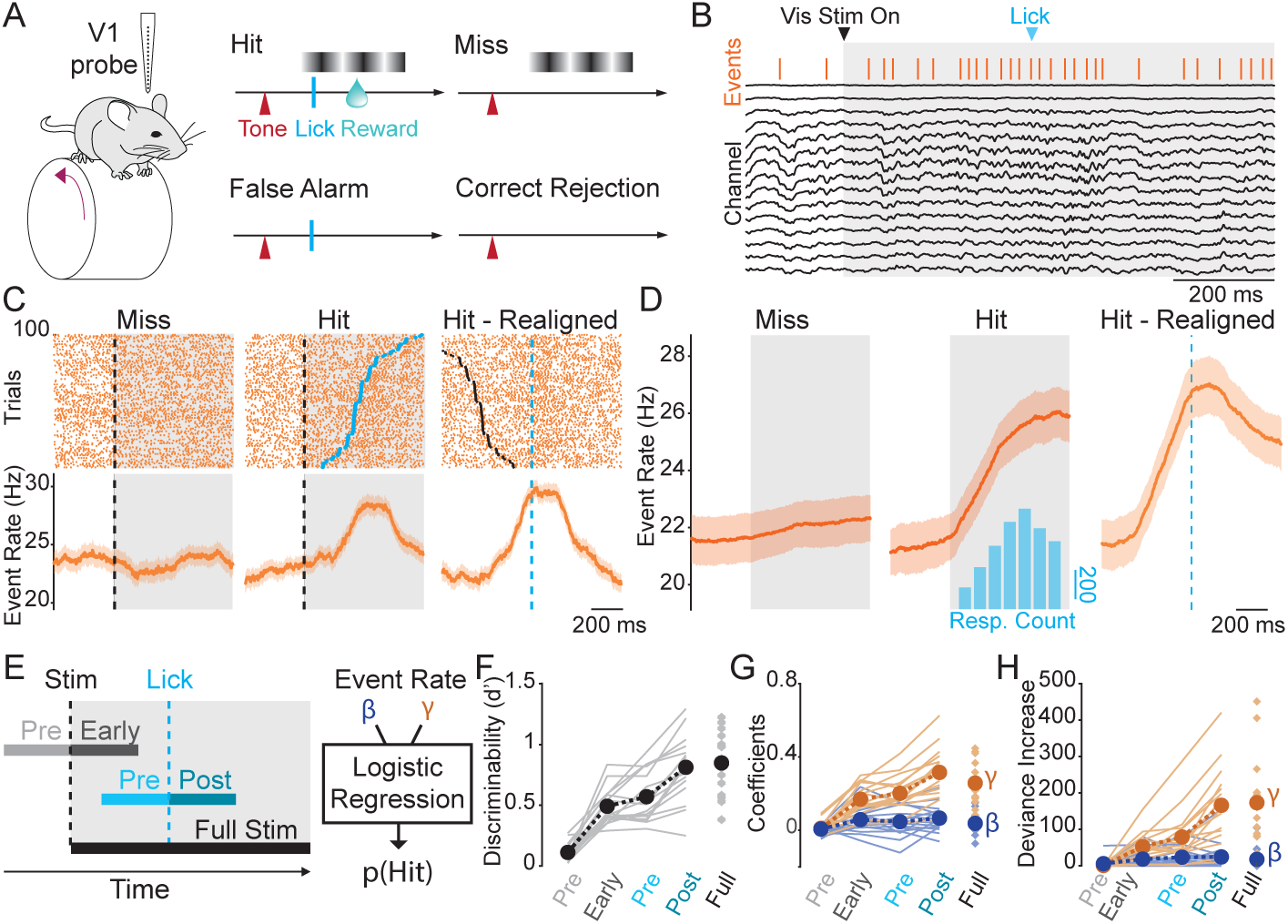
Network events predict behavioral response in a visual detection task. **A**: Schematic of laminar recordings during visual detection task performance. Trial onset is signaled by a tone. If a grating stimulus is displayed, mice can lick to obtain a water reward (Hit). Lick responses made while no stimulus is present on the screen (False Alarm) lead to a time-out. Absence of response to stimulus presentation (Miss) or outside stimulus presentation (Correct Rejection) produce no outcome. **B**: Example recording showing LFP, γ events (orange bars), visual stimulus (gray) and correct lick response (blue arrow) during one trial. **C**: Raster plots of γ event occurrence on 100 randomly selected trials (upper) and average event rate across trials (lower) aligned to stimulus onset (<7.5% contrast; black dotted line) during miss (left) and hit trials (center), and to lick response time (blue dotted line) on hit trials (right). **D**: Population average γ event rate during task trials (n = 16 mice). **E**: Schematic of analysis windows for logistic regression of trial outcome (Pre-Stim: 300ms before stimulus onset, Early-Stim: 300ms after stimulus onset, Pre-Response: 300ms before response, Post-Response: 300ms after response, Full-Stim: Full visual stimulation). **F**: The sensitivity (d’) of the regression increases before response time and is highest right after the response (n = 16 mice). **G**: Model coefficients for γ (orange) and β (blue) events. **H**: Deviance increase upon parameter removal for γ and β events (n = 16 mice). Error bars: mean ± s.e.m. For detailed statistics see Supplemental Table 1.

Increased γ rate prior to behavioral responses could be associated with obtaining a reward. We therefore trained naïve mice to collect free rewards while viewing a gray screen. We observed no significant increase in γ rate leading up to lick responses regardless of reward outcome (Fig. 4), suggesting that γ does not encode generic motor responses or reward signals. To examine whether γ events instead represent a learned association between visual stimulus conditions and reward, we moved the mice to a new paradigm where reward was given exclusively when the lick response occurred during visual stimulation. In this paradigm, γ event rate selectively increased leading up to responses to visual stimuli (Fig. 4, S16). This modulation was rapid, appearing on the first day of the visual paradigm, and independent of behavioral state. Mice were then switched back to the free reward paradigm, leading to immediate loss of the association between γ and behavior (Fig. 4).

**Figure 4.**
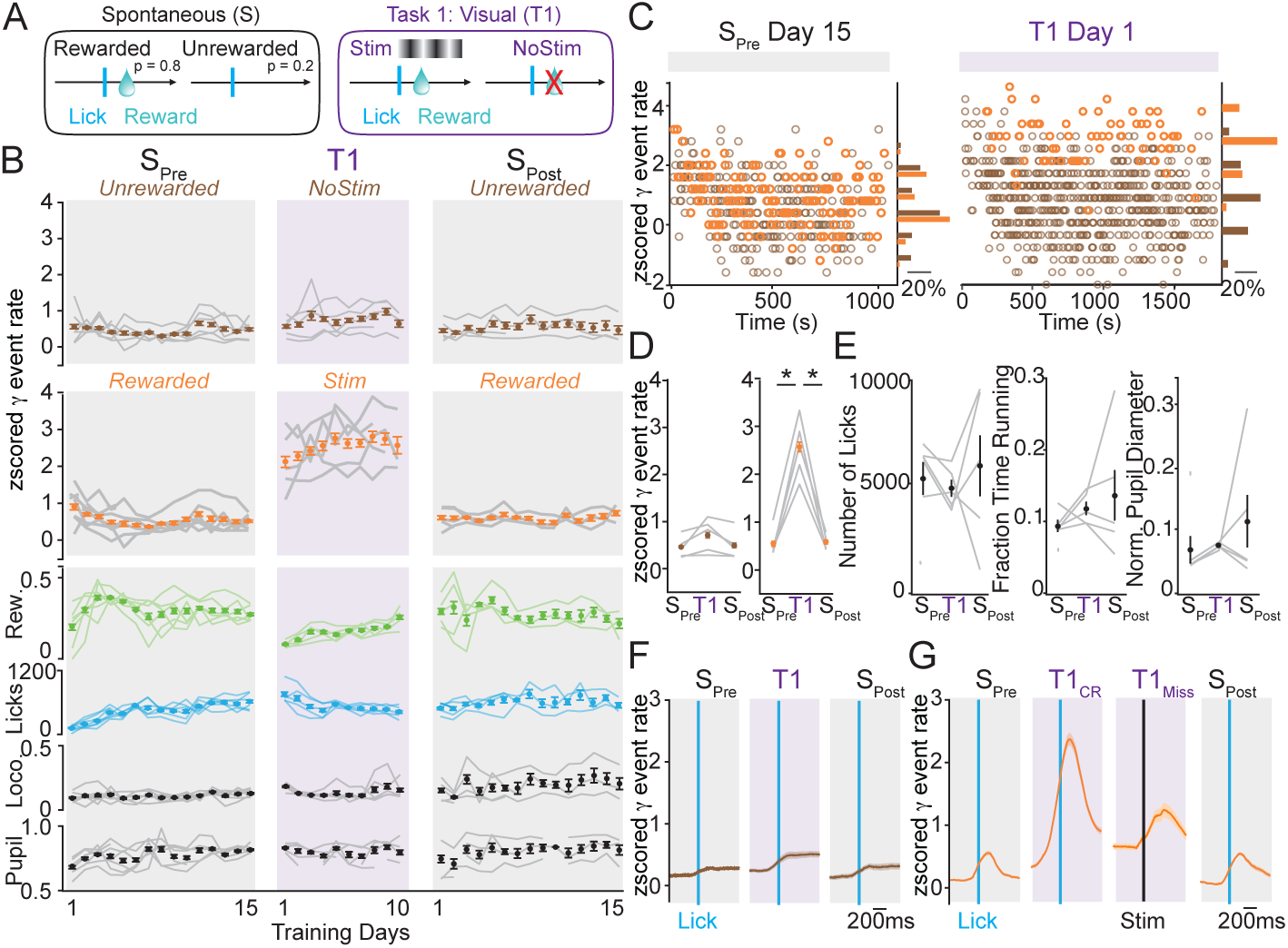
Rapid modulation of network events with changes in task context. **A**: Schematics of trial types for Spontaneous (S) reward paradigm and Task 1 (T1). Mice were first trained to obtain reward freely for 15 days (S_Pre_), then switched to T1, where rewards can only be collected during visual stimuli, for 10 days. Finally, mice were switched back to free rewards (S_Post_) for 15 days. **B**: Normalized γ event rate around rewarded (orange) and unrewarded (brown) responses. Proportion of rewarded trials (green), number of total licks per session (blue), proportion of time spent running (black), and pupil diameter (black) are shown below for each training day. **C:** γ event rate on each unrewarded (brown) and rewarded (orange) trial on day 15 of S_Pre_ and day 1 of T1 in an example mouse. **D:** Overall γ event rate at rewarded (orange) and unrewarded responses during S_Pre_, T1, and S_Post_ paradigms. (*α ≤ 0.05; paired t-test, n = 7 mice). **E:** Average number of licks per session, fraction of time spent running, and average normalized pupil diameter during S_Pre_, T1, and S_Post_ paradigms (n = 7 mice). **F:** Average γ event rate aligned to unrewarded (brown) lick responses during trials during S_Pre_, T1, and S_Post_ paradigms (n = 7 mice). **G:** Average γ event rate aligned to rewarded (orange) lick responses during trials during S_Pre_, T1, and S_Post_ paradigms (n = 7 mice). T1 correct (T1_CR_) and miss (T1_Miss_) trials are shown aligned to lick response and stimulus onset, respectively. (Shaded area: s.e.m). For detailed statistics see Supplemental Table 1.

High-frequency activity in the γ range is a hallmark of arousal and attention processes^5,27,28^, and could be a nonspecific biomarker of changes in global cortical state. Alternatively, increased γ rate in V1 could be linked specifically to visually guided behavior. We found that forced rewards given automatically in association with visual stimulus presentation elicited only a modest increase in γ rate modulation (Fig. S17 A-C). We further examined whether γ responses during task performance were contingent on stimulus modality by recording in V1 during performance of an auditory detection task. In contrast to the visual task, we observed no increased γ rate leading up to correct responses to auditory stimuli (Fig. S17 D-E). Overall, these results suggest that increased γ predictive of performance is modality-specific and sensitive to task context, occurring in V1 when visual information is used to guide behavioral output.

Our findings highlight a novel analytical approach to examine patterned neural activity, linking activity in specific frequency bands to high-resolution, event-based analysis. This approach provides unique insight into the relationship between distinct spatiotemporal patterns of cortical activity, such as β and γ, and perceptual behavior. Using this approach, we were able for the first time to precisely track the rate of individual γ events during performance of a visual task. γ, but not β, rate showed a sharp increase selectively leading up to correct behavioral responses. Furthermore, the relationship between γ events and behavior was rapidly modulated by task context and modality-specific. Additional studies are required to identify network events in other behaviorally relevant frequency bands and to examine the spatiotemporal relationships of network events across cortical areas^29,30^. Our results build upon previous findings in primate^27,28, 31–33^ and rodent^18,34–36^ models and open new avenues to elucidate the functional dynamics of awake cortical activity.

## Author Contributions

QP and JAC designed the experiments. QP and AF developed and validated the CBASS method. QP, AA, JB, and RM collected the data. QP analyzed the data. QP and JAC wrote the manuscript.

## Acknowledgements

The authors thank Drs. Michael J. Higley, Daeyeol Lee, Natalie Schaworonkow and Bradley Voytek for extensive discussions of the data and manuscript and all members of the Cardin and Higley laboratories for helpful input throughout all stages of this study. This work was supported by funding from the NIH (EY022951 and MH113852 to JAC, EY026878 to the Yale Vision Core), a McKnight Scholar Award (to JAC), an award from the Kavli Institute of Neuroscience (to JAC), and a BBRF Young Investigator Grant (to QP).

## Data Availability Statement

The full datasets generated and analyzed in this study are available from the corresponding authors on reasonable request.

## Code Availability Statement

Custom written MATLAB and Python scripts used in this study are available at https://github.com/cardin-higley-lab/CBASS.

## Supplementary Figure Legends

**Figure S1:**
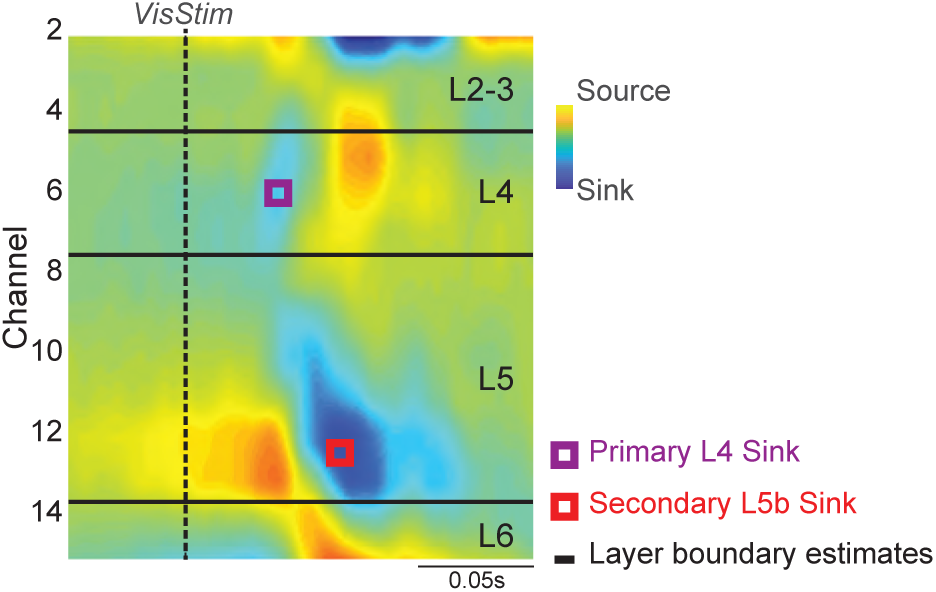
Current source density (CSD)-based mapping of cortical layers. Illustration of the methodology used to estimate the laminar position of LFP channels across cortical layers. The average current source density (CSD) of the response to a high-contrast drifting grating stimulus is computed and consists of a primary sink in cortical layer four (purple) and a secondary sink occurring at longer latencies in layer 5b (red). This allows for a 2-point alignment of a layer boundaries template estimated from histological data (Material & Methods).

**Figure S2:**
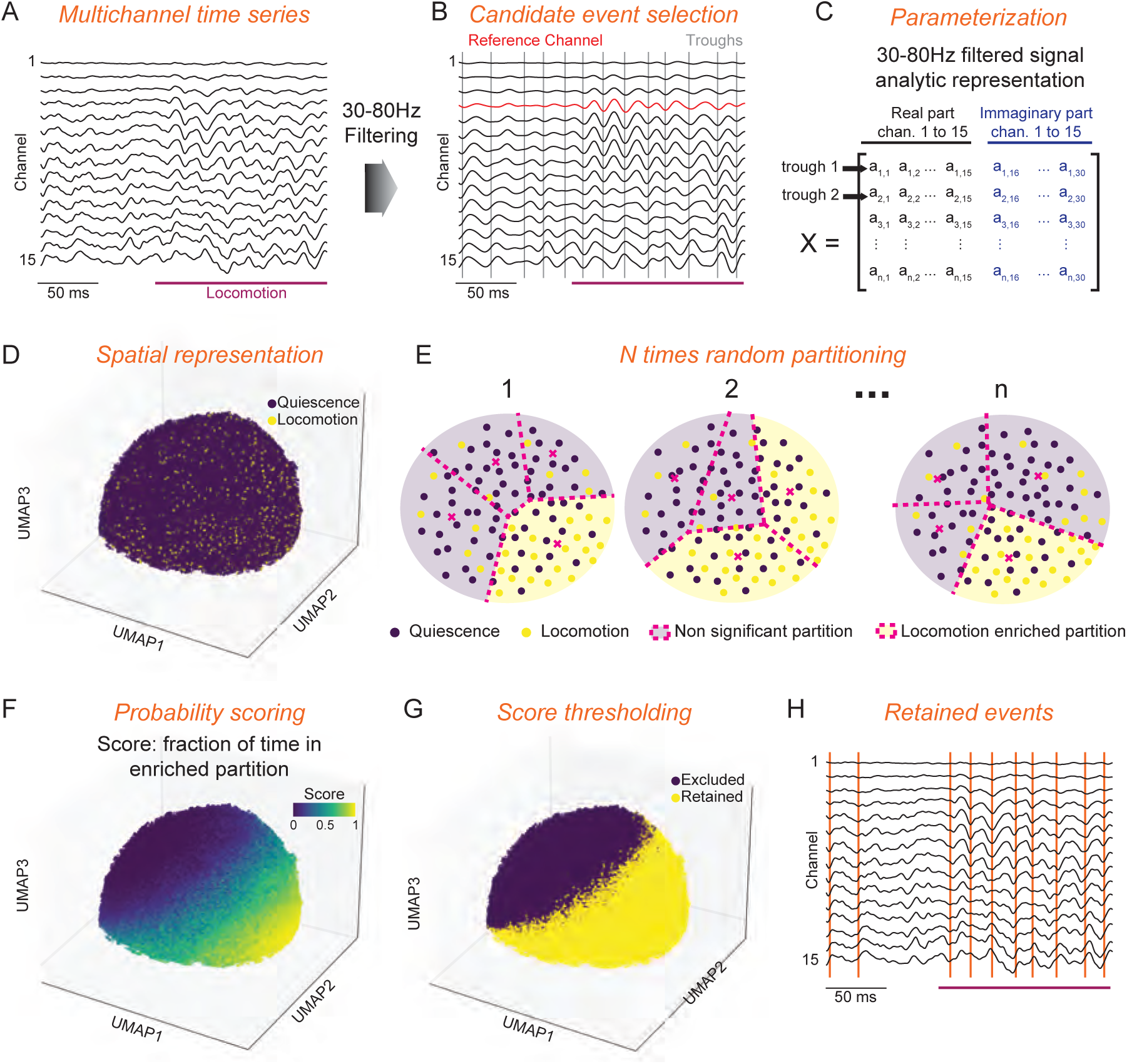
Flow diagram of the CBASS method. CBASS links power increases in a given frequency band during a particular state to events in the temporal domain. As an example, here we look for events responsible for a well-characterized power increase in the gamma range (30-80Hz) in mouse V1 cortex during locomotion. **A**: CBASS starts with a multichannel time series (black) where the state of interest is indexed (i.e. locomotion, purple). **B**: The signal is band-pass filtered in the gamma range. Candidate events (gray bars) are taken at the trough of the filtered signal in a reference channel (red). Here the reference channel is taken as the closest to Layer 4. Different choices of reference channels produce qualitatively similar results but with a shifted event phase (Supplementary Methods). **C**: Spectrotemporal dynamics at the time of each candidate event are parameterized using the real and imaginary part of the analytical representation of the filtered multichannel time series. **D**: Three dimensional UMAP embedding showing the cloud of candidate events in the parametric space. Events occurring during locomotion (yellow) are seemingly present in all regions of the cloud. **E**: CBASS estimates whether specific spectrotemporal activity profiles (i.e. regions of the cloud) occur preferentially during locomotion. The cloud is partitioned randomly, and a binomial test is performed in each partition to test if the occurrence of locomotion is higher than overall. This operation is repeated n times. **F**: An enrichment score is derived for each candidate event as the fraction of time it fell into an enriched partition. This score is stronger in regions of the cloud (i.e. spectrotemporal profiles) associated with a stronger occurrence of locomotion. **G**: CBASS finds the threshold of the enrichment score that produces the most significant separation in the parametric space. **H**: Events whose enrichment score is above the threshold are retained (orange) and noted in the raw data from panel A.

**Figure S3:**
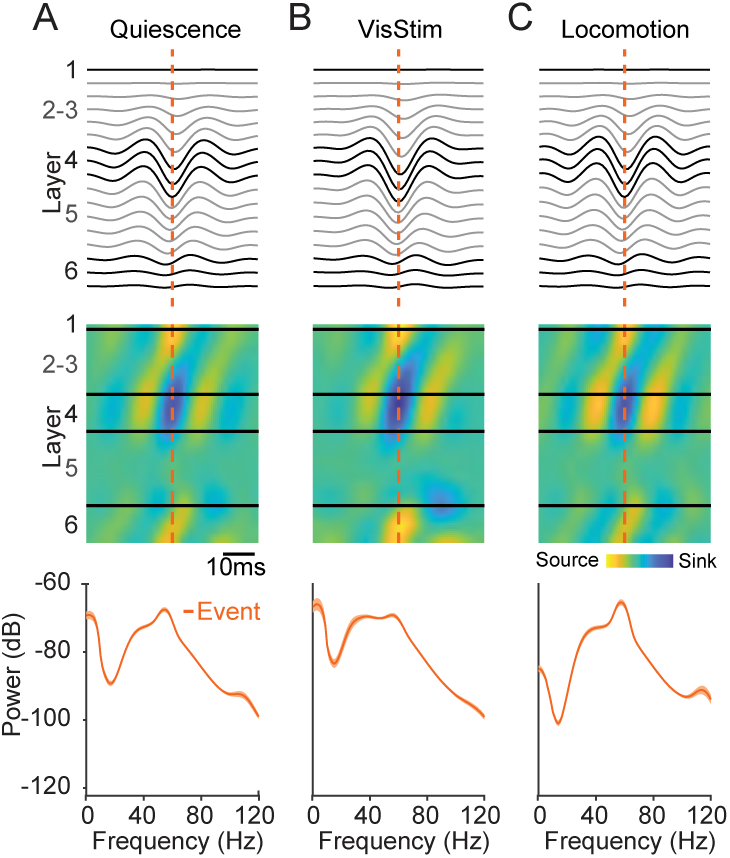
The profile of γ events remains consistent across behavioral states. **A**: Average field potential around γ events (Upper), associated CSD activity (Middle), and power spectrum of the average event field (orange) during quiescence. **B:** Same, during high contrast visual stimulation. **C:** Same, during locomotion.

**Figure S4:**
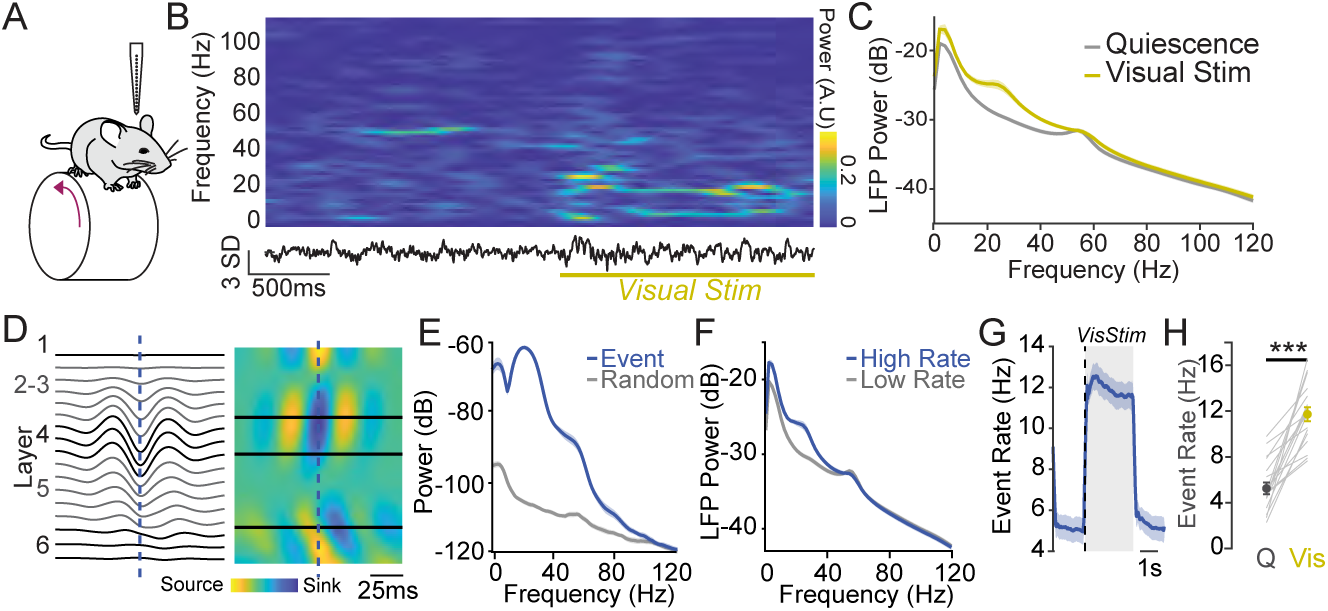
1. CBASS links V1 β power increase during visual stimulation to defined network events. **A**: Mice are head-fixed on a wheel and V1 activity is recorded across cortical layers with 16-channel silicon probes. **B**: Example data showing the LFP in one channel and its short-term Fourier transform during the presentation of a high contrast visual stimulus (yellow). **C**: Average LFP power across channels during quiescence and visual stimulation (19 mice). Visual stimuli evoke increased power in the β range (15-30Hz). **D**: Average field potential around β events and associated CSD activity. Events are associated with an activation of layers 2-3 and 4 followed by an activation of deep layers. **E**: Power of the average LFP event in D (blue) compared to matched random averages (gray). **F**: Power of the LFP when β event rate is high (upper quintile; blue) and when it is low (lower four quintiles; gray). **G**: Event rate around visual stimulation onset (Shaded area: mean ± s.e.m., n = 19 mice). **J**: β event rate increases during visual stimulation (paired t-test, n = 19 mice). For detailed statistics see Supplemental Table 1.

**Figure S5:**
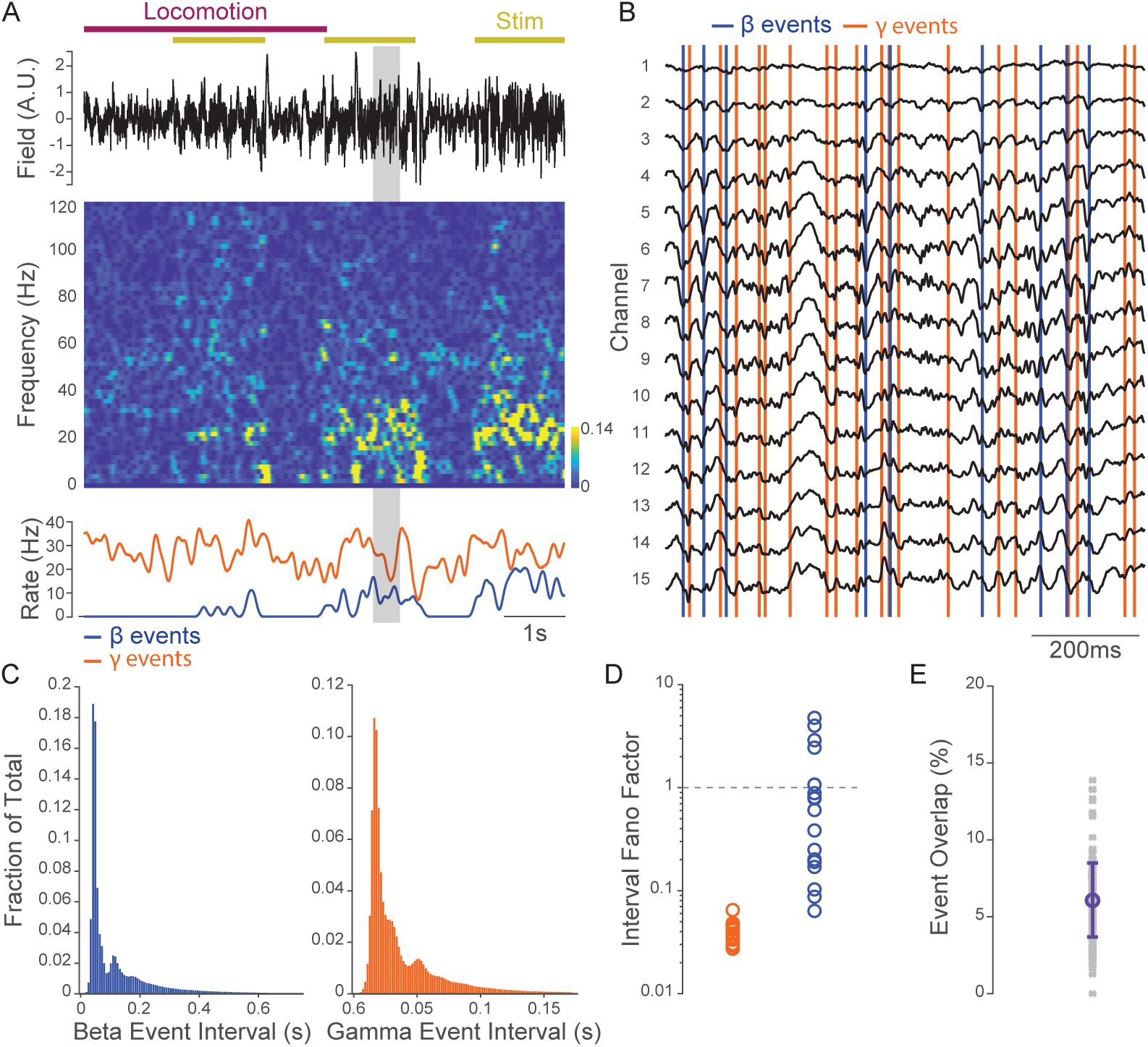
γ and β events identified by CBASS represent distinct processes. **A**: Example recording showing the local field potential in an arbitrary channel in layer 4 during epochs of locomotion and visual stimulation (Upper), its short time Fourier transform (Middle), and the rate of γ (orange) and β events (blue) within a 500ms gaussian sliding window (Lower). **B**: Enlarged version of the gray shaded epoch in panel A showing the LFP in all channels together with detected γ and β events. Event types coincide with distinct dynamics and rarely overlap. **C**: Histograms of the average distribution of the inter-event interval of β (left, blue) and γ (right, orange; n = 19 mice) events. **D**: Fano factor of the inter-event interval distribution of γ and β events (n = 19 mice). γ and β events in most mice have sub-poisson dynamics, indicating that they tend to occur at regularly spaced intervals. **E**: Percent overlap between γ and β events (Gray: mice, Purple: mean ± s.d., n = 201 sessions in 19 mice). For detailed statistics see Supplemental Table 1.

**Figure S6:**
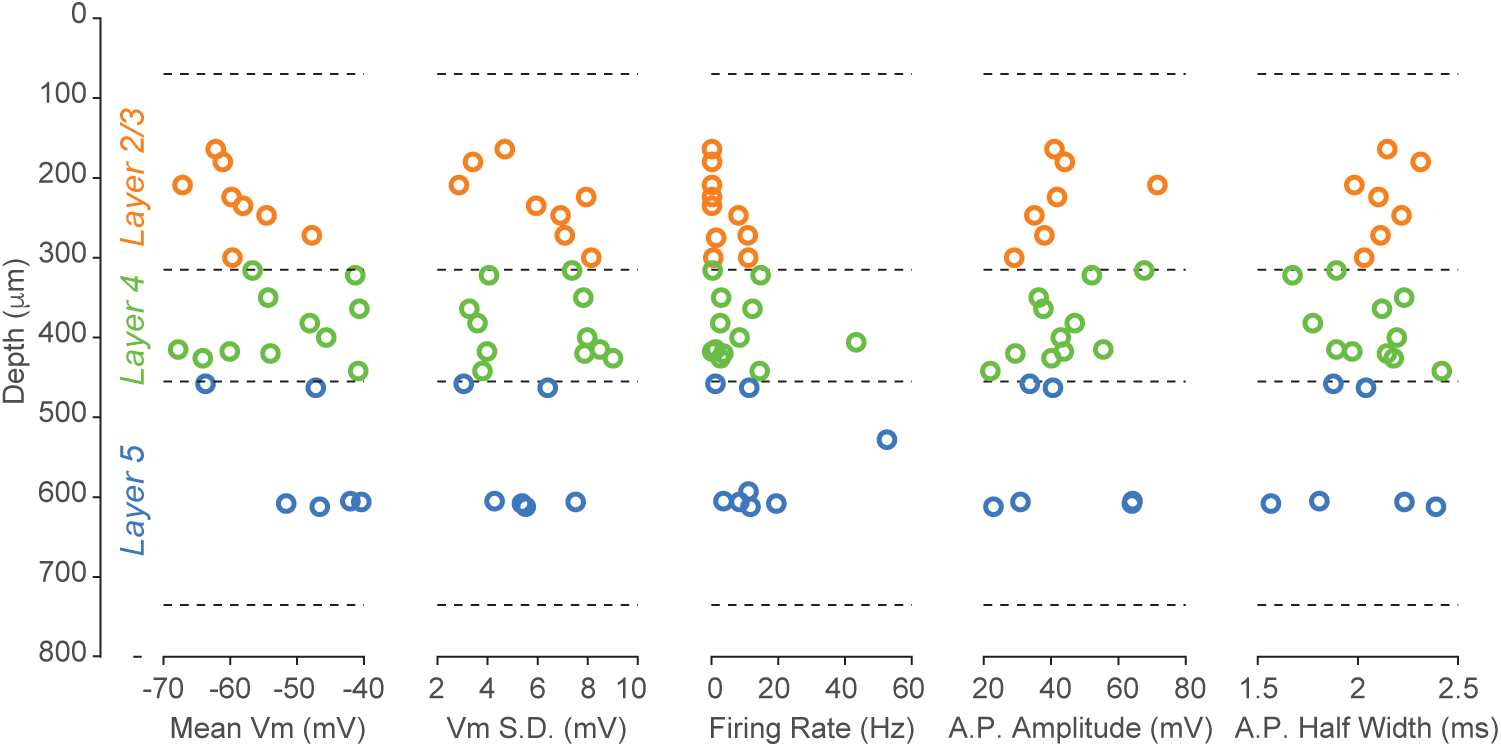
Intrinsic properties and laminar distribution of neurons recorded using *in vivo* whole-cell and cell-attached patch-clamp recordings. From left to right: mean membrane potential (Vm), standard deviation of the membrane potential (VmSD), firing rate, action potential amplitude, and action potential half-width of neurons plotted against recording depth. Neurons between 70 and 315µm were assigned to layers 2-3 (orange, 8 whole-cell, 2 cell-attached), those between 315 and 455 µm to layer 4 (green, 11 whole-cell, 1 cell-attached) and those between 455 and 735 µm to layer 5 (blue, 6 whole-cell, 2 cell-attached). Cell attached recordings were only used to quantify firing rate. One cell in layer 2-3 did not fire any spontaneous action potentials and was only used to quantify Vm activity.

**Figure S7:**
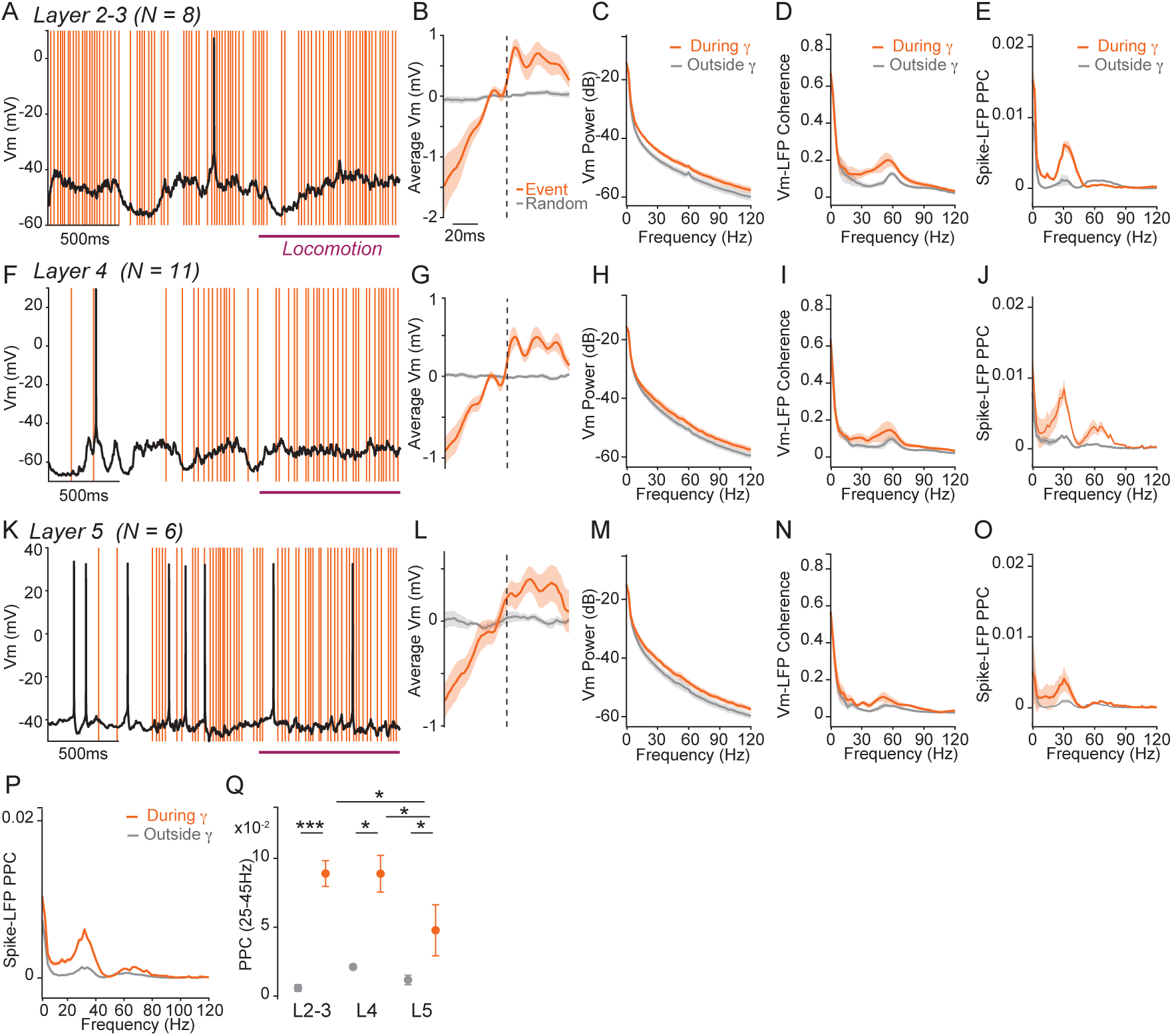
Membrane potential and firing synchronization around γ events. **A**: Example recording of a layer 2-3 neuron with γ events during transition from quiescence to locomotion (purple). **B**: Average membrane potential of layer 2-3 neurons around γ events (orange) and around randomly selected time points (gray) (n = 8 whole-cell recordings). **C**: Power spectrum of the membrane potential of layer 2-3 neurons during (orange) or outside (gray) γ events. Gamma events coincide with an increase of the membrane potential power distributed across the frequency spectrum (n = 8 whole-cell recordings). **D**: Vm-LFP coherence spectra for layer 2-3 neurons during (orange) and outside (gray) γ event cycles, showing selective enhancement of coherence in the γ range (n = 8 whole-cell recordings). **E**: Spike-LFP synchrony spectra for layers 2-3 neurons during (orange) and outside (gray) γ event cycles. Spike-LFP synchrony is quantified using the Pairwise Phase Consistency (Method) and increases during γ event cycles (n = 11 whole-cell/cell-attached recordings). **F**, **G**, **H,** and **I**: Same as A, B, C and D for layer 4 (n = 11 whole-cell recordings). **J**: Same as E for layer 4 (n = 12 whole-cell/cell-attached recordings). **K**, **L**, **M**, and **N**: Same as A, B, C and D for layer 5 (n = 6 whole-cell recordings). **O**: Same as E for layer 5 (n = 8 whole-cell/cell-attached recordings). **P**: Same as E for neurons pooled across layers 2-3, 4 and 5 (n = 30 whole-cell/cell-attached recordings). **Q**: Overall spike-LFP synchrony in the 20-45Hz range, during (orange) and outside (gray) γ event cycles for whole-cell and cell attached recordings in layers 2-3, 4 and 5. Synchrony is enhanced during gamma events and is strongest in Layer 2-3 and Layer 4 neurons (*α = 0.05, **α = 0.01, ***α = 0.001; Welch t-test). Shaded areas and error bars: mean ± s.e.m. For detailed statistics see Supplemental Table 1.

**Figure S8:**
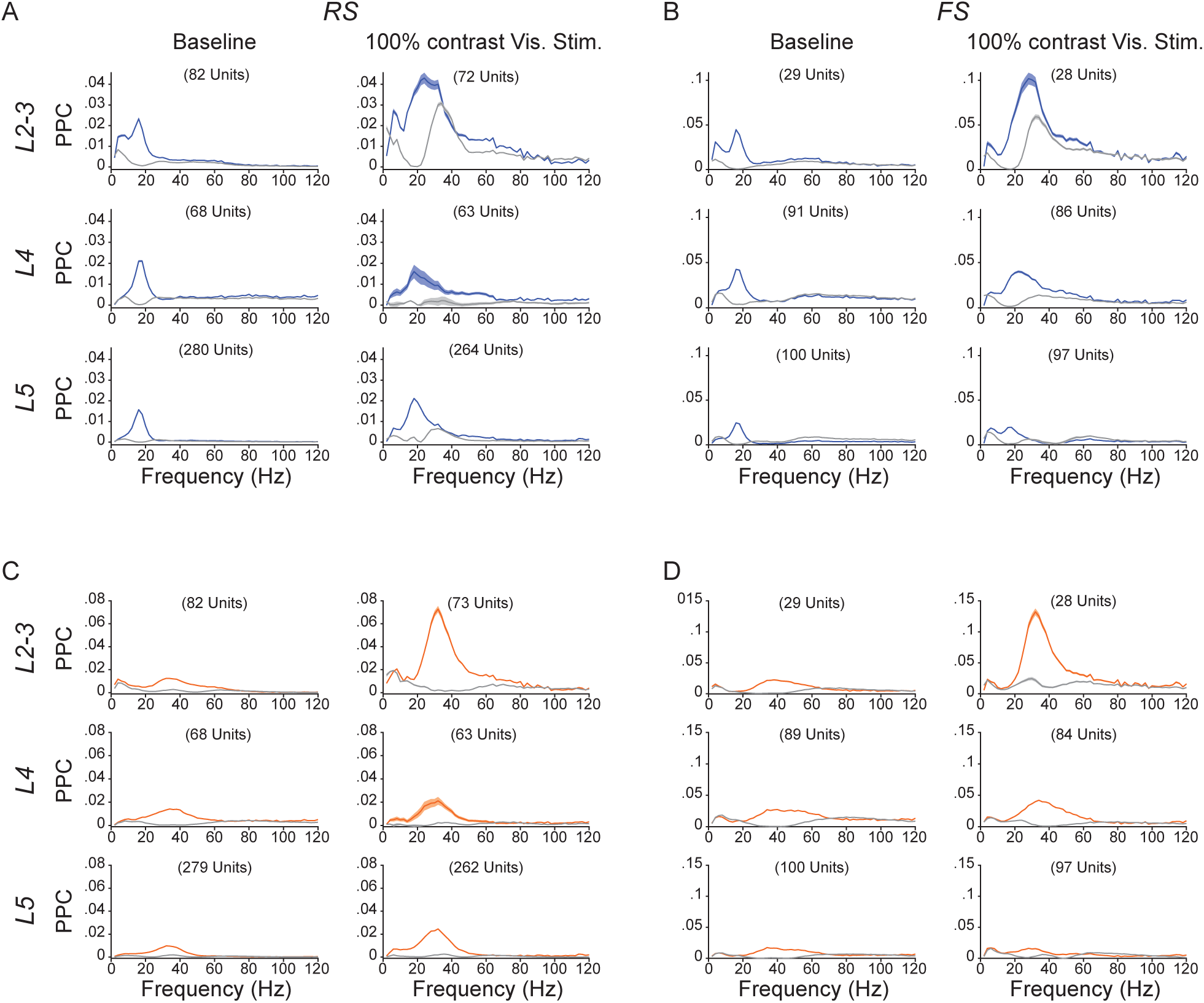
Spike-LFP PPC of RS and FS units. **A**: Spike-LFP synchrony spectra for RS units in layers 2-3 (Upper), 4 (Middle) and 5 (Lower) during (blue) and outside (gray) β event cycles, during baseline (left) and high contrast visual stimulation (right). Spike-LFP synchrony is quantified with the Pairwise Phase Consistency (Method) and increases during β event cycles. **B**: Same as A for FS units. **C** and **D:** Same as A and B during (orange) and outside (gray) γ event cycles. Spike LFP synchrony of RS and FS units increases during γ events. Synchrony in γ events cycles is strongest during visual stimulation for layers 2-3 FS and RS units. For detailed statistics see Supplemental Table 1.

**Figure S9:**
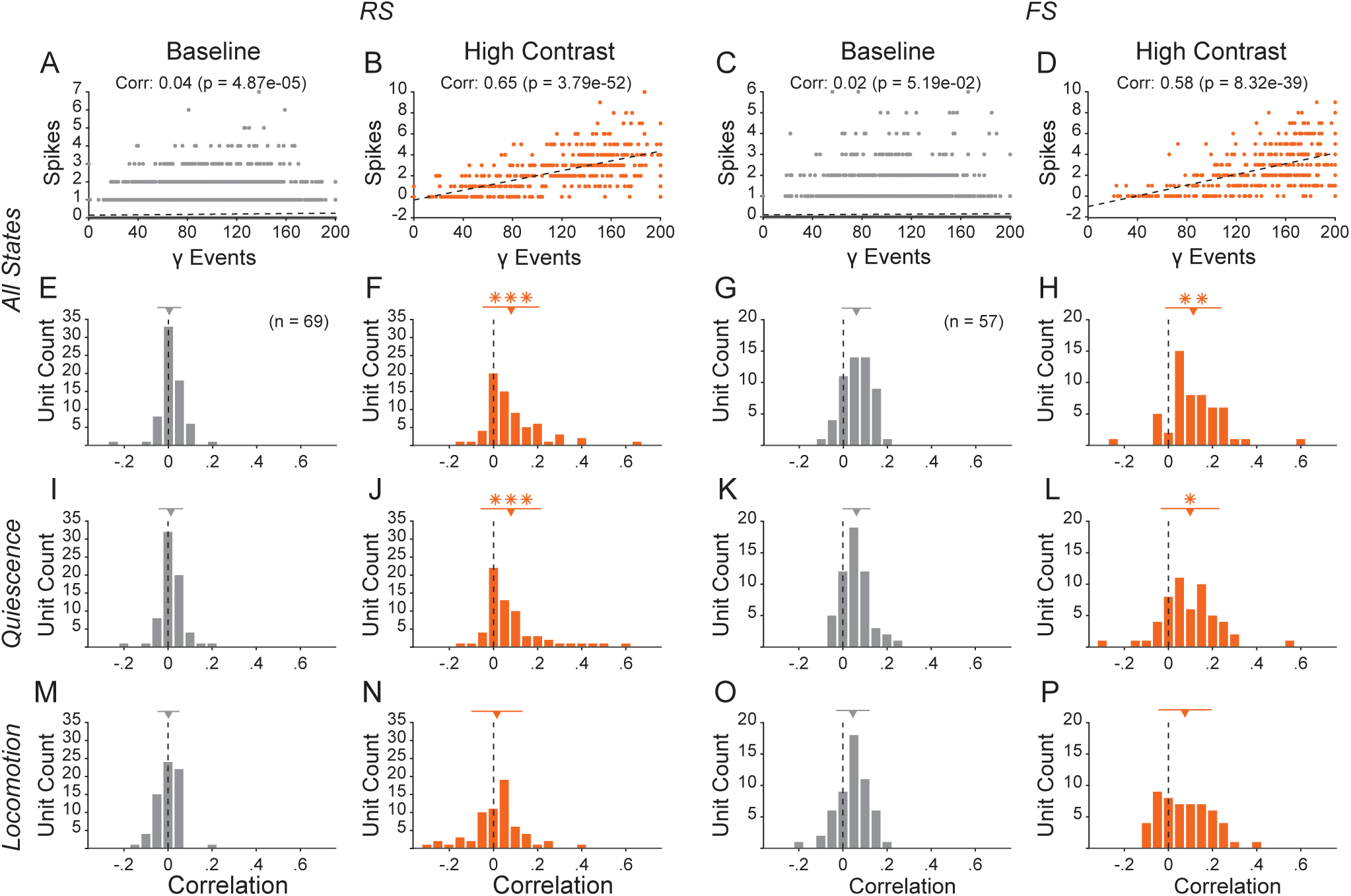
RS unit firing correlates with γ event rates specifically during visual stimuli. **A**: Raster plot of the number of spikes generated by an example RS unit against the number of γ events within 8713 200ms LFP segments recorded during spontaneous baseline activity. **B**: Raster plot of the number of spikes generated by the same example unit against the number of γ events occurring in each of the 425 LFP segments recorded during high contrast visual stimulation. The spike count is correlated with the number of y events during visual stimulation but not during baseline activity. **C**: and **D**: Same as panels A and B for an example FS unit (Baseline: 9577 segments, Stimulation: 429 segments). **E**: Histogram of the correlation values between spike count and γ event number during baseline for 59 RS units (Downward triangle and bars at the top: mean ± S.D.). **F**: Histogram of the correlation values between spike count and γ event number during high contrast visual stimulation for the same units as in panel E, showing a significant increase during high contrast visual stimulation (Downward triangle and bars at the top: mean ± S.D.; *, ** and, *** indicate statistically significant deviation from the mean at baseline with p < 0.05, p < 0.01 and, p < 0.001 respectively; paired t-test). **G** and **H**: Same as E and F for 57 FS units. **I**, **J**, **K**, and **L:** same as E, F, G and H for LFP segments occurring specifically during quiescence. **M**, **N**, **O**, and **P:** same as E, F, G and H for LFP segments occurring specifically during locomotion. For detailed statistics see Supplemental Table 1.

**Figure S10:**
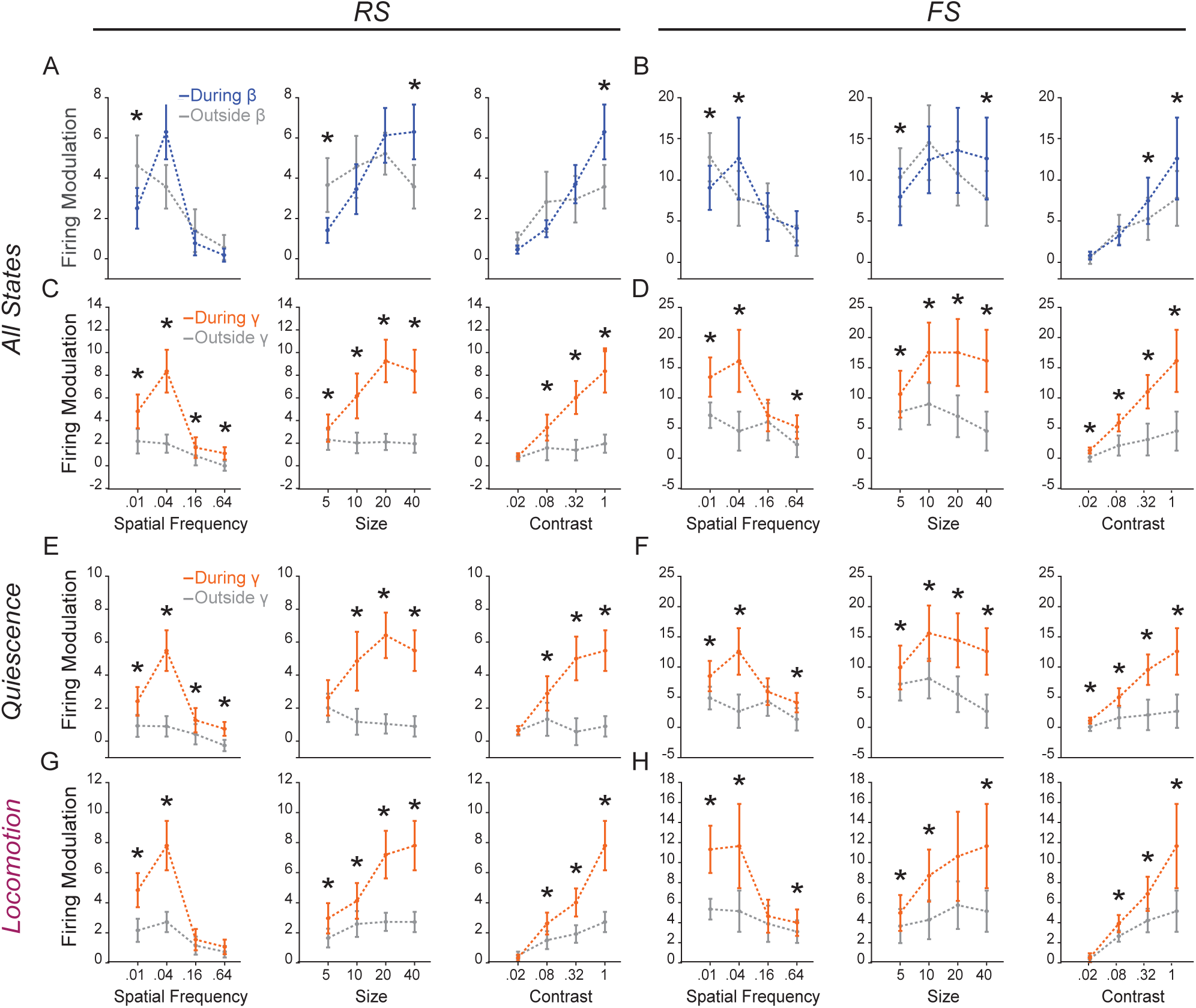
The spike response of RS and FS units to visual stimulation occurs preferentially during γ events. **A**: Modulation of the firing of RS units by gratings of varying spatial frequency (Left), size (Center) and contrast (Right) within (blue) and outside (gray) β event cycles (n = 47 units). Unless otherwise noted, stimuli had a 0.04 cycle/degree spatial frequency, a 40-degree radius and were shown at 100% contrast (*indicates statistically significant difference between modulation within and outside event cycles with p < 0.05; paired t-test). **B**: Same as A for FS units (n = 31 units). Visual feature selectivity was not strongly affected by β events. **C** and **D**: same as A and B for γ events. Firing modulation of RS and FS unit by visual stimuli was markedly stronger during γ events. **E**. and **F**. same as C and D exclusively during epochs of quiescence. **E**, **F**: same as C and D exclusively during epochs of locomotion. Firing modulation by visual stimulation was stronger within γ event cycles across both behavioral states. For detailed statistics see Supplemental Table 1.

**Figure S11:**
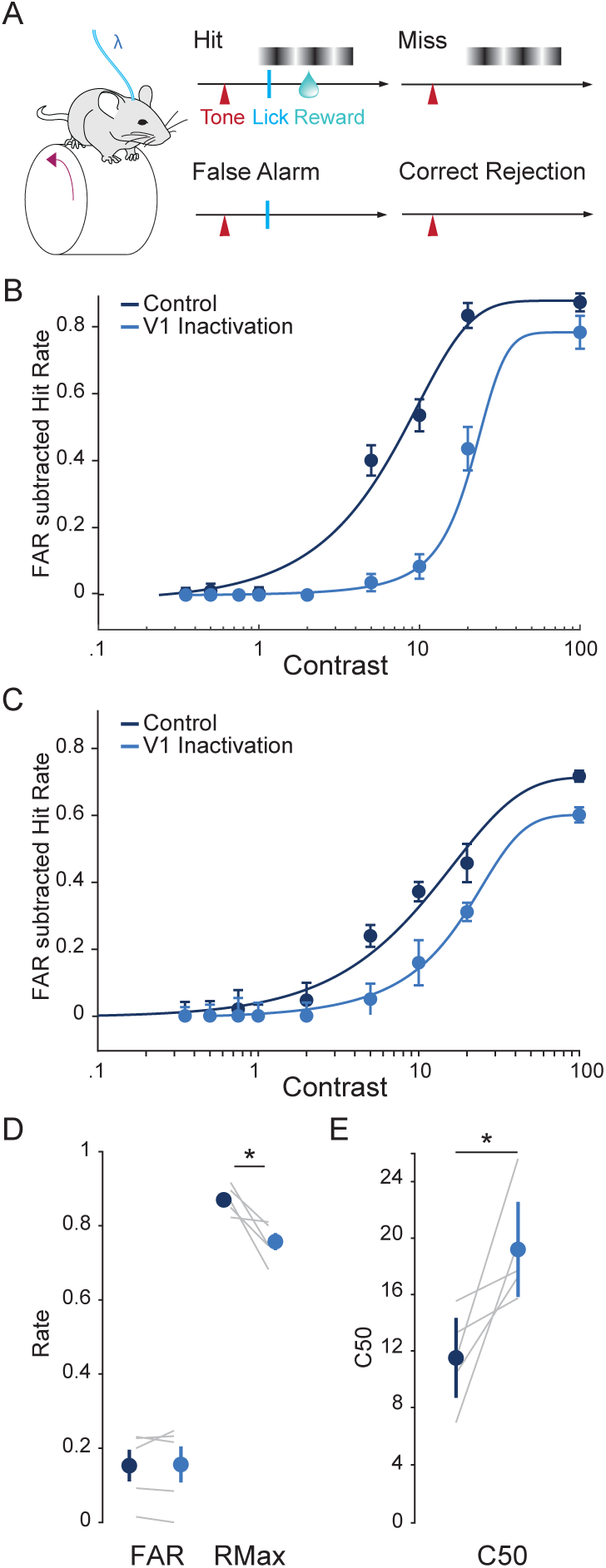
V1 inactivation reduces performance in a visual contrast detection task. **A**: Head-fixed PV-Cre^+/0^ mice injected with a AAV5-ef1α-DIO-ChR2-eYFP virus in V1 (see Supplementary Methods) performed a visual detection task as in Figure 3. On a randomly interleaved subset of trials, blue light was delivered via an optical fiber, bilaterally inactivating V1 through the activation of PV interneurons. **B**: False alarm subtracted hit rate (mean ± s.e.m.) as a function of stimulus contrast during regular trials (dark blue) and during V1 inactivation (light blue) in an example mouse. A sigmoid function is fitted to the hit rate in each condition. **C**: Population average false alarm subtracted hit rate (mean ± s.e.m.) as function of stimulus contrast (n = 5 mice). V1 inactivation reduced detection performance. **D**: False alarm rate (FAR) and hit rate at maximum contrast (RMax) (dark blue) and during V1 inactivation (light blue). V1 inactivation does not affect the FAR but reduces RMax (gray lines: mice; error bars: mean ± s.e.m., *: significant with p < 0.05, paired t-test, n = 5 mice). **E**: Contrast at which the hit rate is 50% (C50) on regular trials (dark blue) and during V1 inactivation (light blue). V1 inactivation increases the C50 (gray lines: mice, error bars: mean ± s.e.m., *: significant with p < 0.05, paired t-test, n = 5 mice). For detailed statistics see Supplemental Table 1.

**Figure S12:**
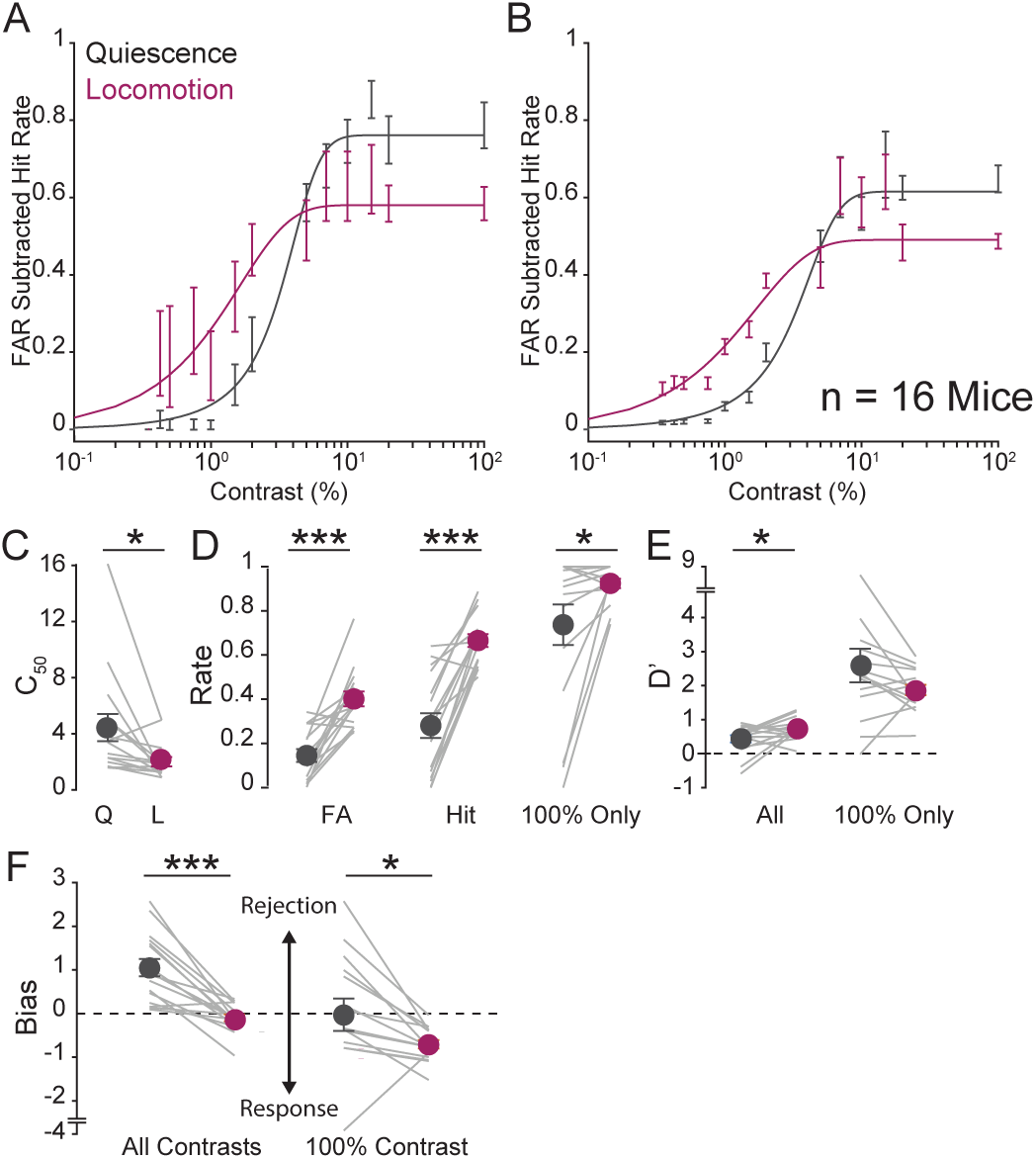
Locomotion enhances visual detection performance and increases bias towards response in a visual contrast detection task. **A**: False alarm subtracted hit rate (mean ± s.e.m.) as a function of stimulus contrast during quiescence (gray) and locomotion (purple) in an example mouse. The hit rate is fitted with a sigmoid curve. **B**: Population average false alarm subtracted hit rate (mean ± s.e.m.) as a function of stimulus contrast during quiescence (gray) and locomotion (purple) (n = 16 mice). **C**: Contrast yielding 50% chance of response (C50) during quiescence (gray) and locomotion (purple). Locomotion is accompanied with a decreased C50 (gray lines: mice, error bars: mean ± s.e.m.). **D**: False alarm rate (FAR), hit rate across contrasts and hit rate at full contrast during quiescence (gray) and locomotion (purple). Locomotion is accompanied with increased hit and false alarm rates (gray lines: mice, error bars: mean ± s.e.m.). **E**: Sensitivity (d’) of the response across contrast and at full contrast during quiescence (gray) and locomotion (purple). Locomotion has a small but significant effect on the sensitivity across contrast (gray lines: mice, error bars: mean ± s.e.m.). **F:** Bias of the response across all contrasts and at 100% contrast during quiescence (gray) and locomotion (purple). Locomotion significantly biases behavior towards responses (gray lines: mice, error bars: mean ± s.e.m., *: significant with p < 0.05, paired t-test, n = 16 mice). For detailed statistics see Supplemental Table 1.

**Figure S13:**
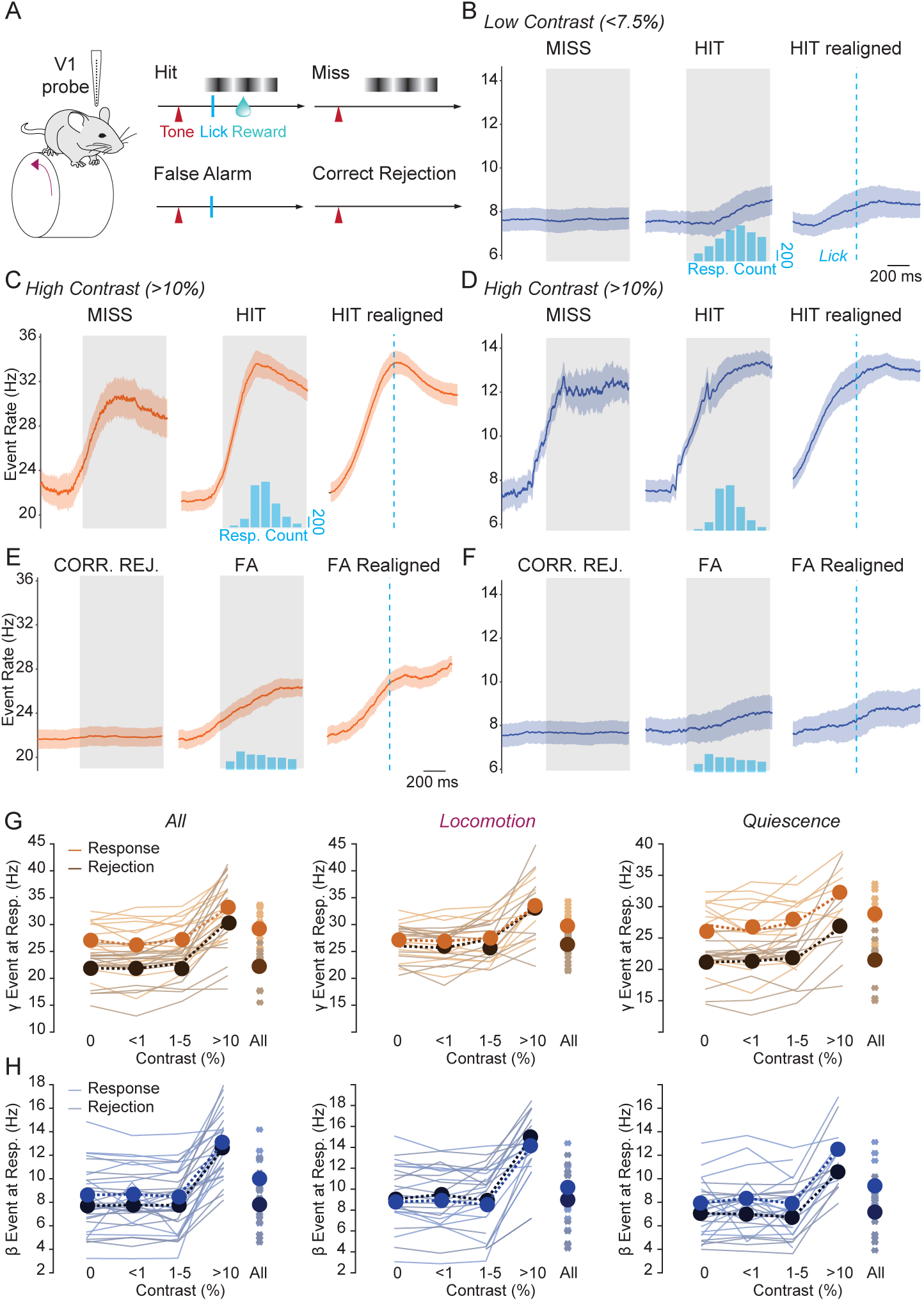
Selective increase in γ, but not β, events prior to behavioral response in a visual detection task. **A**: Head-fixed mice perform a visual contrast detection task while V1 activity is recorded with chronically implanted silicon probes (see Fig. 3). **B**: Average β event rate across 16 mice during low contrast trials (< 7.5%). Event rate is aligned to stimulus onset during miss (left) and hit trials (middle), and to response time on hit trial (complementary to Fig. 3D). β event occurrence is not significantly higher on hit trials (shaded area: mean ± s.e.m., gray box: time of visual stimulus presentation). **C**: Same as panel B for γ events during high contrast (> 10%) trials. **D**: Same as panel C for β events. **E**: γ event rates during 0% contrast (no go) trials. (gray box: time when visual stimulus becomes possible). **F**: Same as panel E for β events. β event occurrence is not significantly higher during FA trials. **G**. Rate of γ event at in the 300ms following response or average response time (rejections) for trials with stimuli of increasing contrasts, across all behavioral states (Left), during locomotion (Center) and during quiescence (Right). The rate of γ events is significantly higher at response than during rejection across contrasts, except during locomotion (thin lines: mice, error bars: mean ± s.e.m.). **H**: Same as panel G for β events. There is no significant difference in β event rate between response and rejection trials. For detailed statistics see Supplemental Table 1.

**Figure S14:**
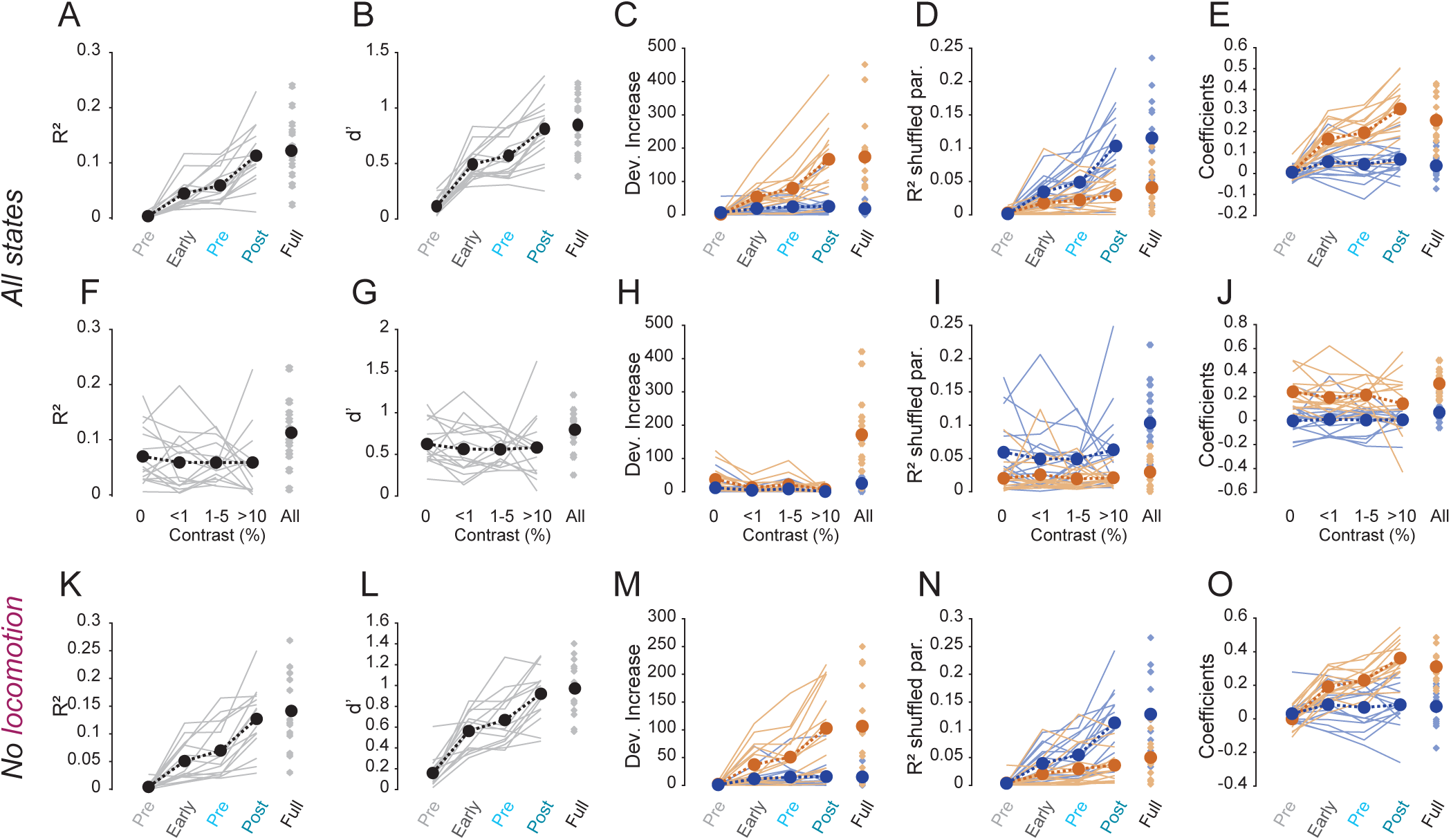
γ event occurrence predicts the trial-by-trial outcome of visual detection task performance across stimulus contrasts and behavioral states. **A**: McFadden’s R-squared (R^2^) of a logistic regression of trial outcome based on γ and β event rate in different windows around stimulus onset, lick response or average response time (for rejection trials) (Pre-Stim: 300ms before stimulus onset, Early-Stim: 300ms after stimulus onset, Pre-Response: 300ms before response, Post-Response: 300ms after response, Full-Stim: Full visual stimulation, thin line: mice, thick dotted line: average across 16 mice). **B**: Same as panel A using the sensitivity (d’) to measure regression performance. Prediction increases though the trial. **C**: Deviance increase upon parameter removal. **D**: R^2^ after parameter shuffling. **E**: Regression coefficients show that γ event occurrence has the strongest influence on model prediction (thin line: mice, thick dotted line: average across 16 mice). **F**, **G**, **H**, **I** and **J**: same as A, B, C, D and E for visual stimulation with increasing contrasts in the Post-Stimulus window. Model performance is stable across contrasts, suggesting that the predictions do not arise simply from contrast-dependent responses in γ or β. **K**, **L**, **M**, **N** and **O**: same as panels A, B, C, D and E, excluding trials where locomotion occurred at any point within 2s of trial onset. Locomotion-related increases in γ event occurrence do not account for model performance. For detailed statistics see Supplemental Table 1.

**Figure S15:**
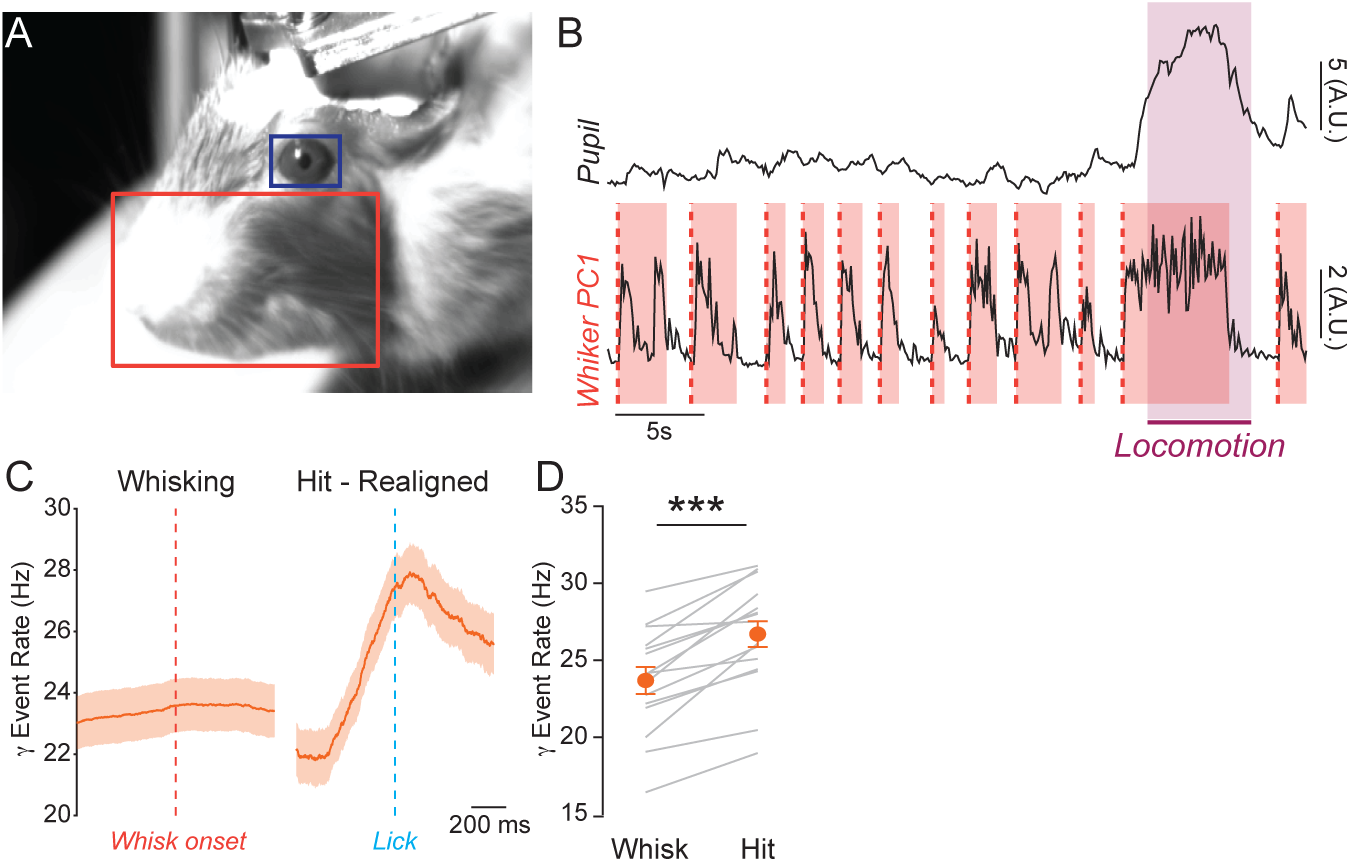
γ event rate is not modulated by whisking. **A**: Images from a facial video recording of a head-fixed mice running on a wheel while V1 activity was monitored with chronically implanted silicon probes. Areas of interest are defined for the pupil (blue) and the whisker pad (red). **B:** Excerpt showing pupil diameter and first principal component of whisker pad pixel-value variance around a locomotion bout (purple box). Whisking epochs (red boxes) were defined with a change point algorithm (Supplementary Methods). **C**: Average γ event rate across 16 mice around whisking onset (red bar, left) and lick response (blue bar) during low contrast trials (< 7.5%, right). **D:** Event rate is higher during correct response trials than after whisking onset (t-test; p < .001, n = 17 mice). For detailed statistics see Supplemental Table 1.

**Figure S16:**
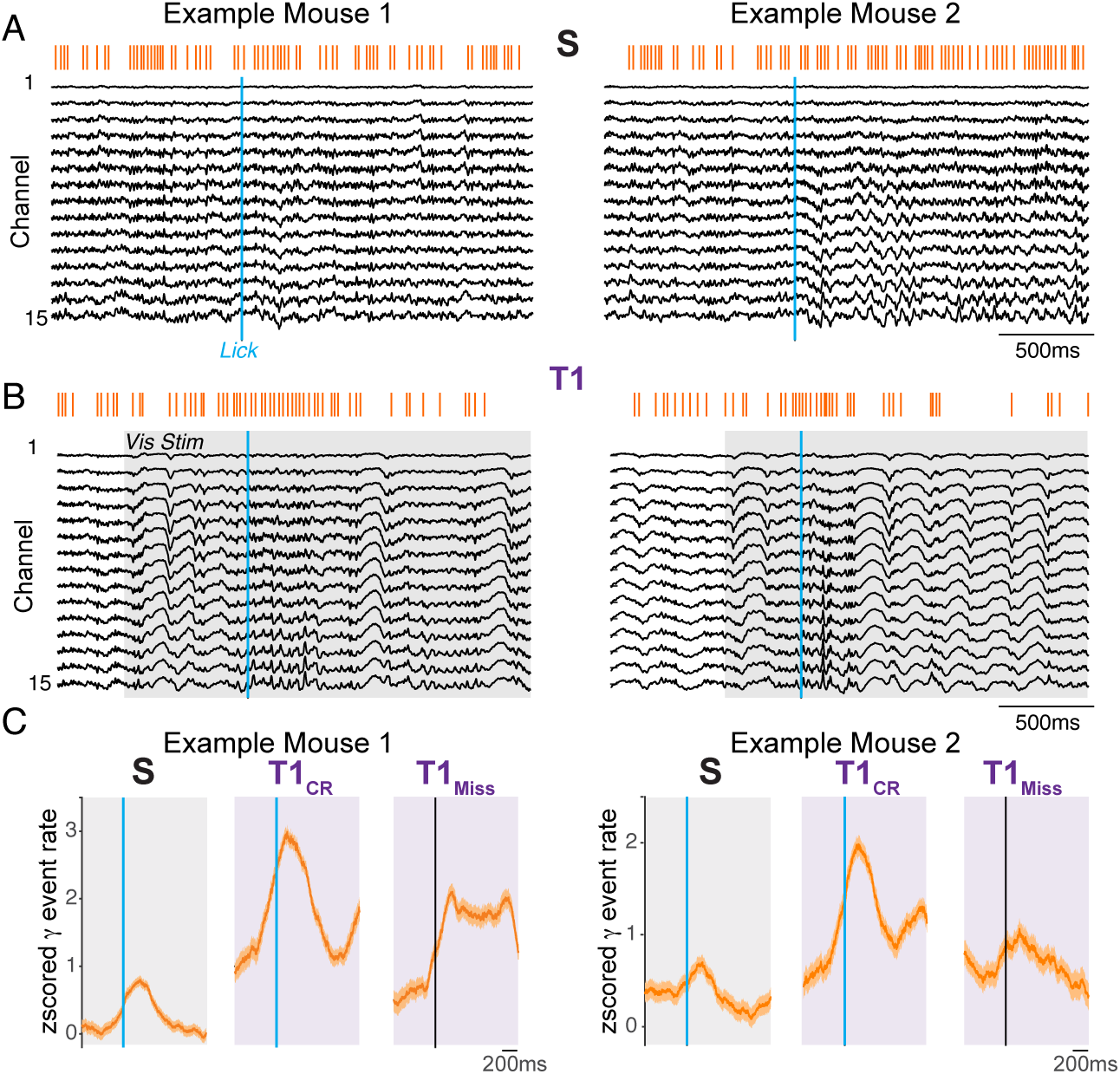
γ event modulation around spontaneous and visually cued rewarded lick responses in example mice. Data from V1 recordings of 2 example mice during task performance as in Figure 4. **A:** Multichannel LFP and γ events (orange) around example rewarded lick responses on the last day of the spontaneous paradigm (S_Pre_) where mice can freely lick for reward (Fig. 4A). **B:** Example multichannel LFP and γ events around rewarded lick responses on the first day of the Task 1 (T1) paradigm where rewards are only distributed during visual stimuli (gray square). **C:** Average normalized γ event rate around rewarded lick-responses (blue) and stimulus onset for miss trials (black) in the S_Pre_ and T1 paradigms. For each example mouse, visually cued responses elicit a stronger increase in γ event rate than does visual stimulation alone.

**Figure S17:**
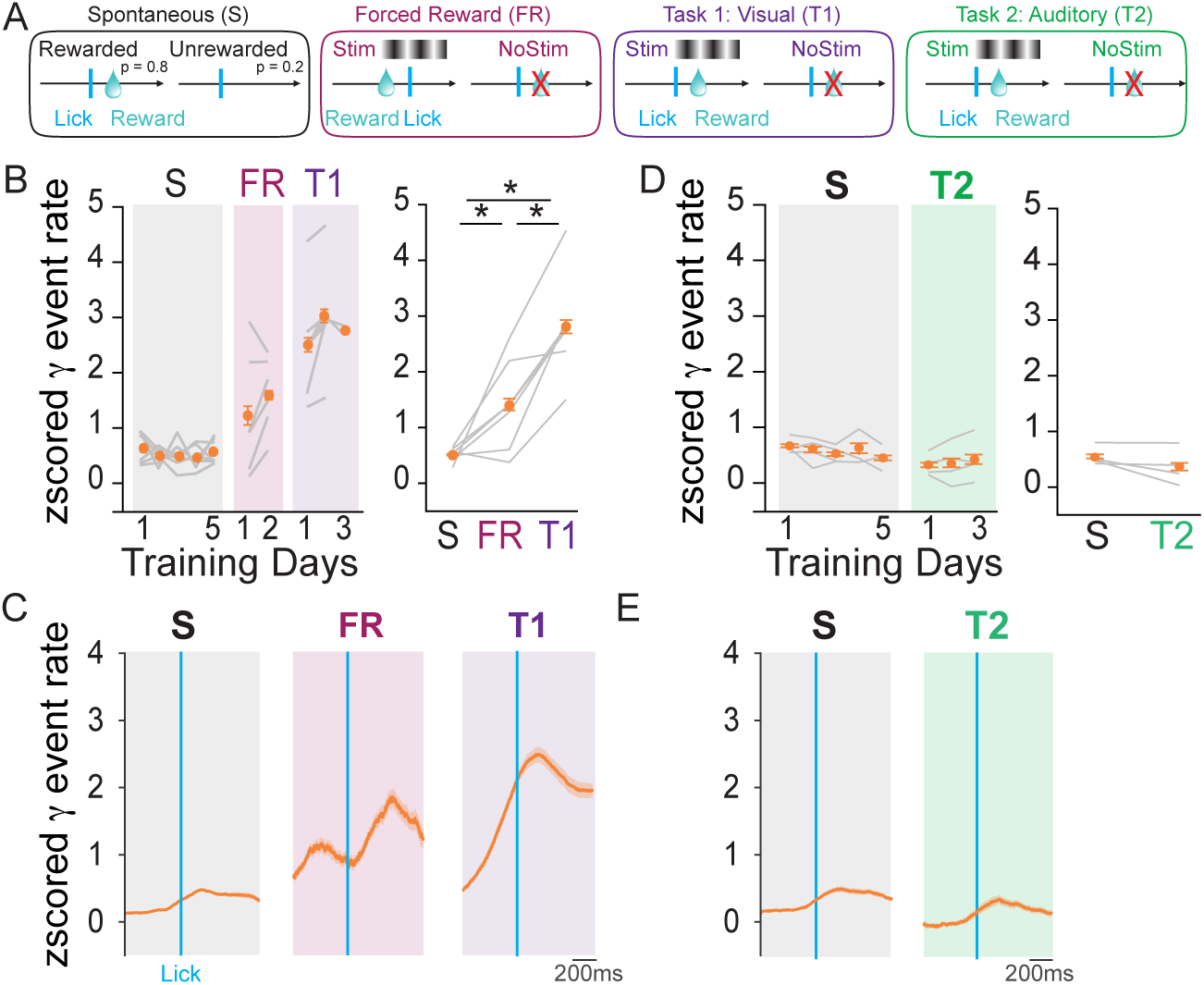
Rate of γ event occurrence around response across behavioral paradigms. **A**: Schematics of trial types for behavioral paradigms using Spontaneous licking with free reward (S), forced reward (FR), visual detection (T1), and auditory detection (T2). **B:** Mice were trained on the spontaneous reward (S) task for 5 days and then switched to the forced reward task (FR) where reward is automatically delivered at the onset of each visual stimulus for 2 days. Mice were then trained on the visual detection task (T1) where reward is delivered only when a lick response occurs during the visual stimulus. Normalized γ event rate (orange) around rewarded responses is shown for S, FR, and T1 training days (n = 4 mice). **C:** Normalized γ event rate (orange) around rewarded responses is shown for S, FR, and T1 training days (n = 4 mice). **D:** Mice were trained on the Spontaneous paradigm for 5 days, then on T2 for 3 days. Left: Normalized γ event rate for rewarded responses across training days. Right: Overall γ event rate for rewarded responses over S and T2 (n= 4 mice). γ event rate in V1 does not increase during auditory guided responses. **E:** Normalized γ event rate around rewarded responses (blue) in the S, and T2 paradigms. For detailed statistics see Supplemental Table 1.

## Material & Methods

### Animals

Male and female C57Bl/6 mice were kept on a 12h light/dark cycle, provided with food and water ad libitum, and housed individually following headpost implants. A subset of mice used for optogenetic experiments were heterozygous for PV-ires-Cre (PV-Cre^+/0^) (strain# 008069, Jackson Laboratory). All animal handling and experiments were performed according to the ethical guidelines of the Institutional Animal Care and Use Committee of the Yale University School of Medicine.

### Surgery

Mice were anesthetized with isoflurane (1.5% in oxygen) and maintained at 37°C for the duration of the surgery. Analgesia was provided with subcutaneous injections of Carpofen (5mg/kg) and Buprenorphine (.05mg/kg). Lidocaine (1% in 0.9% NaCl) was injected under the scalp to provide topical analgesia. Eyes were protected from desiccation with ointment (Puralube). The scalp was resected and the skull cleaned with Betadine. A surgical screw was implanted on the skull between the eyes and nuts were glued to the skull above the bregma suture, allowing the fixation of a headplate with bolts. For chronic electrophysiology, 2 craniotomies were performed respectively above V1 on the left hemisphere (~0.15mm diameter; 2.5mm laterally from lambda) and above the cerebellum (0.4mm diameter; ~2mm posterior to lambda). An A16 probe with a CM16 connector (Neuronexus) was lowered into V1. Ground and reference wires were inserted above the cerebellum. For acute electrophysiology, a circular plastic ring (~2.5mm diameter) was glued on the skull above V1. The skull inside the ring was protected with cyanoacrylate. For optogenetic manipulations, craniotomies were performed above V1 on each hemisphere, 1μl of AAV5-ef1α-DIO-ChR2-eYFP (Addgene) was injected at a depth of 300µm in each hemisphere and optical canulae (Doric Lenses Inc.) were positioned above the dura. Craniotomies were protected with Gelfoam (Pfizer), and all implants were affixed to the skull with dental cement (Metabond, Parkell Industries).

### Electrophysiology

Mice were habituated to handling and head fixation for 3-5 days prior to electrophysiological recordings. For chronic recordings, mice were head-fixed on a wheel (Vinck et al., 2015) and their implants were connected to the recording apparatus (DigitalLynx system, Neuralynx). The most superficial contact point was used as a reference.

For acute silicon probe and patch clamp recordings, two small craniotomies (~0.1mm, <0.1mm apart) were performed above V1 under isoflurane anesthesia. Analgesic was provided as described above and mice were moved back for >2h in their home cage to recover from anesthesia. Mice were head-fixed on the wheel. The ring situated above V1 was filled with artificial cerebrospinal fluid (ACSF; in mM: in mM: 135 NaCl, 5 KCl, 5 HEPES, 1 MgCl_2_, 1.8 CaCl_2_ [adjusted to pH 7.3 with NaOH]), an AgCl reference electrode placed in the bath and an A16 probe (Neuronexus) was lowered into V1. Glass pipettes (4–6 MΩ) were pulled from borosilicate capillaries (Outer diameter: 1.5mm; Inner diameter 0.86; Sutter Instrument) and filled with and internal solution (in mM: 135 potassium gluconate, 4 KCl, 10 HEPES, 10 phosphocreatine, 4 MgATP, 0.3 Na_3_GTP, [adjusted to pH 7.3 with KOH; osmolarity adjusted to 300 mOsmol]). Pipettes were lowered into V1 and whole cell patch clamp configurations were obtained at depth ranging from 164 to 742µm. After achieving intracellular access, a minimum delay of 5 minutes was included before recording to allow cortical activity to recover normal dynamics. Intracellular recordings were amplified with a Multiclamp 700 B amplifier (Molecular Devices). In all experiments, pupil (Vinck et al., 2015) and facial motion (Stringer et al., 2019) were recorded at 10Hz using an infrared camera (FLIR). Local Field Potentials, wheel motion, and timing signals for face movies, visual stimulus, and behavior were acquired at a 40KHz sampling rate.

### Visual stimulation and behavior hardware

Visual stimuli were generated using the Psychtoolbox Matlab extension (Kleiner et al., 2007) and displayed on a 17’’ by 9.5’’ monitor situated 20cm in front of the animal (Visual Detection task) or 15 cm from the right eye (all other behavioral tasks; passive visual stimulation). Screen display was linearized and maximum luminance was adjusted to ~140 cd.sr/m^2^. An iso-luminant grey background was displayed between visual stimuli. Task-related actions were implemented through sensors and actuators interfaced with a microcontroller (Arduino Due; Teensy 3.2) connected to a computer running custom routines in Matlab. Waterspouts were positioned using a servomotor (Hi-tec). Responses were detected through an optical sensor (Optex-FA) and water delivery was controlled using solenoid valves (Asco). When behavior was performed during electrophysiological recordings, timing signals for spout movement, response, and reward delivery were sent from the microcontroller to analog ports on the DigitalLynx system.

### Visual response measurements

The visual response of single units was tested using vertical gratings drifting leftward with a 1Hz temporal frequency and centered on the receptive field at the recording site. Gratings were presented for 3s and separated by a 2s interstimulus interval. Unit responses properties were investigated at all combinations of 4, 8, 32 and 100% contrasts, 0.01, 0.04, 0.16 and 0.64 cycle/degree spatial frequencies, and 10-, 20-, 40- and 80-degree diameters (64 combinations total).

### Behavioral experiments

For behavioral training, mice were water rationed and maintained between 82% and 88% of their initial weight. Reward consisted of 3μl water droplets. All visual stimuli were full-screen drifting gratings with a spatial frequency 0.04 cycle/deg and temporal frequency of 2Hz and were displayed for 1 second. Auditory stimuli consisted of pure tones at 2KHz. On trials where mice responded by licking, the stimulus was displayed for an additional 2 seconds during reward consumption.

#### Visual detection task

Training was divided into 5 stages. **1.** Mice were first trained to collect water freely from the waterspout. Reward was given at regular intervals. Mice were moved to the next stage when they made 100 responses in a 20-minute session. **2.** Mice were habituated to the trial structure and to associate reward to high-contrast (100%) visual stimuli. The waterspout was moved within reach and after a 4s delay, a pure tone (4kHz, 200ms) signaled the onset of a trial. Visual stimuli were displayed after a randomized interval (0.5 to 1.2s) and a reward was delivered at stimulus onset. Mice could collect an additional reward if they licked during the visual stimulus. The spout was moved out of reach at the end of trial for an additional interval (1.5 to 3.5s). Mice were moved to the next stage after two 30-minute sessions. **3.** Mice had to lick during visual stimulus presentation (100% contrast) to receive a reward. Mice were moved to the next stage when then responded correctly on more than 80% of trials within a 30-minute session. **4.** No-go trials were introduced. Stimuli were omitted after the tone on 30% of trials. If animals made a response when stimuli were not present on the screen, the waterspout was moved away, and mice incurred a 10s timeout. Mice were moved to the next stage when their Hit rate was >80% and their false alarm rate <20%. Sessions lasted 45 minutes. **5.** Contrast was varied to test psychophysical performance. Task structure was otherwise identical to stage 4.

To test the role of V1 in task performance, PV-Cre^+/0^ mice were bilaterally injected with 1µl of AAV5-Ef1a-DIO-Chr2-eYFP viral vector (titer: ~10^12^ vg/mL, Addgene) and implanted with optic canulae (Doric Lenses Inc.) as described above. Mice were trained on the visual detection task until stage 5. After 5 days on stage 5, and no less than 30 days after implantation, V1 was inactivated on 30% of trials with bilateral optogenetic activation of parvalbumin expressing interneurons. Light pulses (55ms, 10Hz) were delivered through an insulated multi-mode optical fiber (200µm diameter, 0.53 NS, Thorlabs) coupled to a 473nm solid state laser (Opto Engine LLC). Laser power was adjusted to produce an output of ~110mW/mm^2^. Pulse timing was controlled through a shutter (Thorlabs). Pulse trains started 300ms before stimulus onset and were maintained until the end of the trial.

To investigate how gamma event rate at response time depended on reward contingencies we used training schedules consisting of combinations of the following paradigms:

#### Spontaneous paradigm

No stimuli were displayed. Mice were given rewards at Poisson-distributed time intervals (λ= 10s) to ensure a flat hazard rate. Lick responses made at any time led to additional rewards with an 80% probability.

#### Task 1 Visual paradigm

Reward were given only when lick responses were made during visual stimuli. Stimuli appeared on the screen at Poisson distributed time intervals (λ= 9s).

#### Task 2 Auditory paradigm

Rewards given were given only when lick responses were made during auditory stimuli. The task structure was otherwise identical to Task 1.

#### Forced reward paradigm

Rewards were passively given at the onset of visual stimuli. An additional reward was given upon reward collection. The task structure was otherwise identical to Task 1.

Training schedules were always initiated with the Spontaneous paradigm in mice having no prior experience in behavioral experiments other than habituation to head-fixation and handling.

### Preprocessing

Data were analyzed in Matlab 2018b (Mathworks) using custom scripts. All time-series were down-sampled to 2KHz (patch clamp recording) or 1KHz (chronic recordings) and aligned. Local field potential (LFP) recordings were high-pass filtered at 1Hz using a 2nd order Bessel filter and z-scored across channels. LFP channels were mapped onto cortical layers using the current source density (CSD) profile of visual responses (Fig. S1). Recordings of membrane potential (Vm) were curated using a custom-made procedure to delineate epochs suitable for processing. Epochs were retained if (1) spike threshold was within −40 +/− 2mV, (2) spike peak was above −20mV, (3) Vm values outside spikes stayed in the [−85 −40] mV range. Junction potentials were not corrected but were estimated as −14.9mV as described previously (Perrenoud et al., 2016). For event-triggered averages of Vm, spikes were removed [−2 to 5] ms from peak and missing values were interpolated with cubic splines. Pupil diameter was measured from movies with a custom procedure (Vinck et al., 2015). Pupil diameter was normalized to the average pupil diameter during locomotion for comparison between recordings. The first principal component of whisker pad motion energy was computed from the same movie using FaceMap (Stringer et al., 2019). Pupil diameter and facial motion were interpolated and aligned to the other time series. Epochs of running and whisking activity were defined using a change point algorithm detecting local changes in the mean and variance of running speed and whisker pad motion (Vinck et al., 2015). Briefly, moving standard deviations of speed and facial motion energy were computed with a defined temporal window. The length *t* of this window determines the temporal resolution of the changepoint analysis and was set to 4s for running speed and 500ms for facial motion. A first estimate of locomotion/whisker motion onset/offset times were then taken as the time when the moving standard deviations exceeded/ fell below 20% of its range above minimum. Estimates were refined in a window *t* around each onset/offset time by computing the time points corresponding to the maximum of the *t*-windowed moving forward/backward Z-score.

### Single unit clustering

Single units were extracted from LFP recording using spikedetekt and clustered using klustakwik2 (Rossant et al., 2016). Cluster were visualized and sorted using the phy-gui (https://github.com/cortex-lab/phy) together with a custom matlab GUI to compute quality metrics. Single-unit clusters were generally retained if less than 0.2% of inter-spike intervals were inferior to 2ms and if their isolation distance and L-Ratio were superior to 15 and inferior to 0.01 respectively (Schmitzer-Torbert et al., 2005). Isolation distance and L-Ratio are biased by spike number so deviations to those rules were occasionally allowed for unit of low firing rate if their waveform was well above noise.

Fast spiking (FS) and regular spiking (RS) units were defined as described previously (Vinck et al., 2015). Briefly, the average normalized waveforms of all units were clustered with the k-means method based on 2 parameters: peak to trough time, and repolarization (i.e. defined as the value of the normalized waveform 0.45ms after peak). FS units had higher repolarization values and shorter peak-to-trough times than RS units.

### CBASS

CBASS (Clustering Band-limited Activity by State and Spectral features) ties a power increase in a defined frequency band (i.e., gamma (30-80Hz)) during a particular state (i.e., running) to the occurrence of defined events in the temporal domain. A detailed description is available in the appendix below and implementations in matlab and python are available on (https://github.com/cardin-higley-lab/CBASS). Briefly, the multichannel LFP is filtered in the band of interest and candidate events are selected at the troughs of the filtered signal in a reference channel (Fig. 1D, Fig. S2A-B). The spectrotemporal dynamics underlying each candidate event are parameterized using the real and imaginary part of the analytical representation (matlab function *Hilbert*) of the filtered signal in each channel (Fig. S2C). Candidate events form a cloud in this parametric space where neighbors have similar spectro-temporal dynamics (Fig. S2D). The event cloud is split randomly into ***n*** partitions and a binomial test is performed in each partition to determine if events happen during the state of interest (i.e. running) at higher frequencies than overall. Partitioning is repeated ***N*** time (Fig. S2E). A state enrichment score is calculated for events as the fraction of time they fell into an enriched partition (Fig. S2F). An optimization procedure is then applied to find the threshold yielding the most significant distance between events having a low and a high enrichment score in the feature space (Fig. S2G). Events above threshold are retained (Fig. S2H). Here we used ***n*** = 20 partitions and ***N*** = 1000. Different settings for these parameters have only a marginal influence on the result of the procedure.

### Layer alignment of LFP and CSD across recordings

To compute the average field potential around CBASS events across recordings, the LFP was linearly interpolated across channels to a common grid of laminar position (Fig. 1F, Fig. S4D). The CSD was derived as the second spatial derivative of the LFP across interpolated laminar positions.

### Comparison of network activity within and outside CBASS event cycles

CBASS events are aligned to the trough of the band-pass filtered LFP in a reference channel. We defined each event’s boundaries as the peaks surrounding the event’s trough. Peak and troughs were determined as the 0 and π valued time points of the argument (matlab function *abs*) of the analytic representation (matlab function *hilbert*). Activity inside the event boundaries thus fell within a cycle centered on the trough. Epochs during and outside all CBASS event cycles were pooled separately and compared.

### Spike distribution around CBASS events

Spike distribution around CBASS events was computed as follows. For a selected unit, the lag separating each spike from the nearest CBASS events was estimated. A histogram of lag values was then computed and normalized by total spike count. Histograms were averaged across units.

### Event rate normalization

Normalized rates for CBASS-detected events were calculated as follows. A baseline event rate ***p*** was computed over samples. The variance of the rate over a window of ***n*** samples was estimated assuming a binomial distribution as 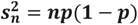. The normalized rate of events over a window of ***n*** samples was then taken as 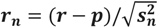 where ***r*** is the event rate over samples and can be thought of as the number of standard deviations away from baseline.

### Unit firing modulation by visual stimulation

Modulation of single-unit action potential firing by visual stimulation was calculated similarly to normalized event rate. A baseline firing rate ***r*** was computed over samples outside visual stimuli. The variance of the rate over a window of ***n*** samples was estimated assuming a binomial distribution as 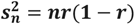. The modulation of event firing for each stimulus modality samples was then taken as 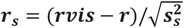 where ***rvis*** is the visually evoked firing rate and ***s*** is the number of samples within the visual stimulation period. Firing modulation can be thought of as the number of standard deviations away from the mean baseline rate. The baseline firing rate of each unit was computed separately within and outside CBASS event cycles.

### Spectral analysis

The spectral power of a given time series was derived with Welch’s method. Each channel was divided into 500ms overlapping segments (75% overlap). Each segment was multiplied by a Hamming window and their Fourier transform was computed (matlab function *fft*). Power was derived as 10 times log10 of the squared magnitude of the Fourier Transform and expressed in dB. Power was averaged over segment and channels.

The spectral power of event-triggered averages was derived with a minimum bias multi-taper estimate (Riedel and Sidorenko, 1995). This differs from a classical multi-taper estimate in that Slepian tapers are replaced by a sinusoidal tapers sequence defined as:

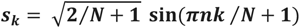

where ***N*** is the number of samples in the triggered average, ***n*** is the sample number and ***k*** is the order of the taper. Sinusoidal tapers produce a spectral concentration almost comparable to that achieved with a Slepian sequence while markedly reducing local bias. The number of tapers was chosen to yield a bandwidth of .8Hz following the formula: *K* = *round*((*4 πNB* / *r*) – 1) where ***B*** is the bandwidth and ***r*** is the sample rate. Triggered averages were multiplied by each taper. Spectral power was then computed as described above and averaged over tapers.

For coherence and spike phase locking estimation, spectro-temporal representations were first derived either for a set of frequencies using a wavelet transform (matlab function *cwt*) and a Morlet wavelet (matlab identifier *cmor1-2*) or across a full frequency band by computing the analytical representation of the filtered signal (matlab function *Hilbert*). Coherence was defined as:

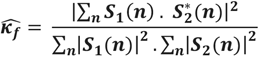

where ***S_k_(n)*** is the spectro-temporal representation of signal ***k*** for sample ***n*** at the frequency ***f***. ***κ_f_*** has a positive bias of ***(1 – κ_f_)/*N** where ***N*** is the number of samples. The bias was subtracted from the estimate. Spike phase locking was estimated using the Pairwise Phase Consistency (Vinck et al., 2010) defined as:

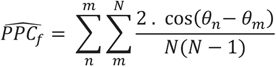

where ***θ_k_*** is the phase of the signal for frequency ***f*** at the time of spike ***k*** and ***N*** is the total number of spikes. PPC provides an unbiased estimate of spike phase locking. However, estimate can be noisy if the spike number is inferior to 250. Thus, population estimates of PPC were derived by pooling spikes from all selected neurons and the variance over neurons was estimated with a leave-one-out Jackknife procedure (Shao and Wu, 1989).

### Logistic regression

Logistic regressions of trial outcome in our visual detection task were performed using the matlab function *glmfit* and a logit transfer function. Logistic regression models return an estimated of the probability of response for each trial. The log-likelihood of regression models was calculated by summing the log-likelihood of each trial’s outcome given the probabilities returned by the model and assuming a Bernoulli distribution. Model performances were tested using likelihood ratio tests and quantified with McFadden’s R-Squared and a sensitivity metric (d’). McFadden’s R-Squared was defined as:

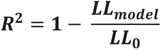

where ***LL_model_*** represents the log-likelihood of the regression and ***LL_0_*** represent the log likelihood of the null model (i.e. the likelihood of the data assuming that all trials have an equal probability of success corresponding to the mean hit rate). Sensitivity was defined as

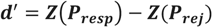

where **Z** is the inverse standard normal distribution and ***P_resp_*** and ***P_rej_*** represent the average probability of response returned by the model for response and rejection trials respectively. The impact of each regressors was assessed in two ways. 1. Regression was recomputed 1000 times after shuffling regressor’s values over trials. A p-value for the significance of each regressor’s impact was derived as the percentage of R-Squared on shuffled values superior to the actual R-Squared of the model. 2. Regression models was compared to a model where each regressor was taken away and the significance of the regressor’s contribution was estimated with a likelihood ratio test. The magnitude of a regressor’s contribution was measured using the increase in deviance. Deviance represents the difference of predictive power from a saturated model giving a perfect prediction (i.e. the likelihood of each trial is 1). It is defined as:

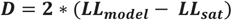

where ***LL_model_*** is the log-likelihood of the model and ***LL_sat_*** is the log-likelihood of the saturated model. Significance was estimated separately for each mouse. Statistical significance across mice was assessed by pooling p-values using Fisher’s method.

### Statistics

Statistics in each figure panel are described in Supplementary Table 1. Except where otherwise noted, tests were performed using mice as the statistical unit. When indicated independent p-values derived on individual mice were pooled using Fisher’s method. Multiple comparisons were corrected using Benjamini-Yukutieli’s procedure for false discovery rate (Benjamini and Yekutieli, 2001).

## Appendix - CBASS: Detailed Methods

This documentation is also available on (https://github.com/cardin-higley-lab/CBASS/wiki)

LFP power often increases in a specific frequency band during specific events or behavioral states. For example, in the visual cortex of mouse, beta (15-30Hz) increases during visual stimulation while gamma (30-80Hz) increases during running (Fig. 1).

**Figure 1.**
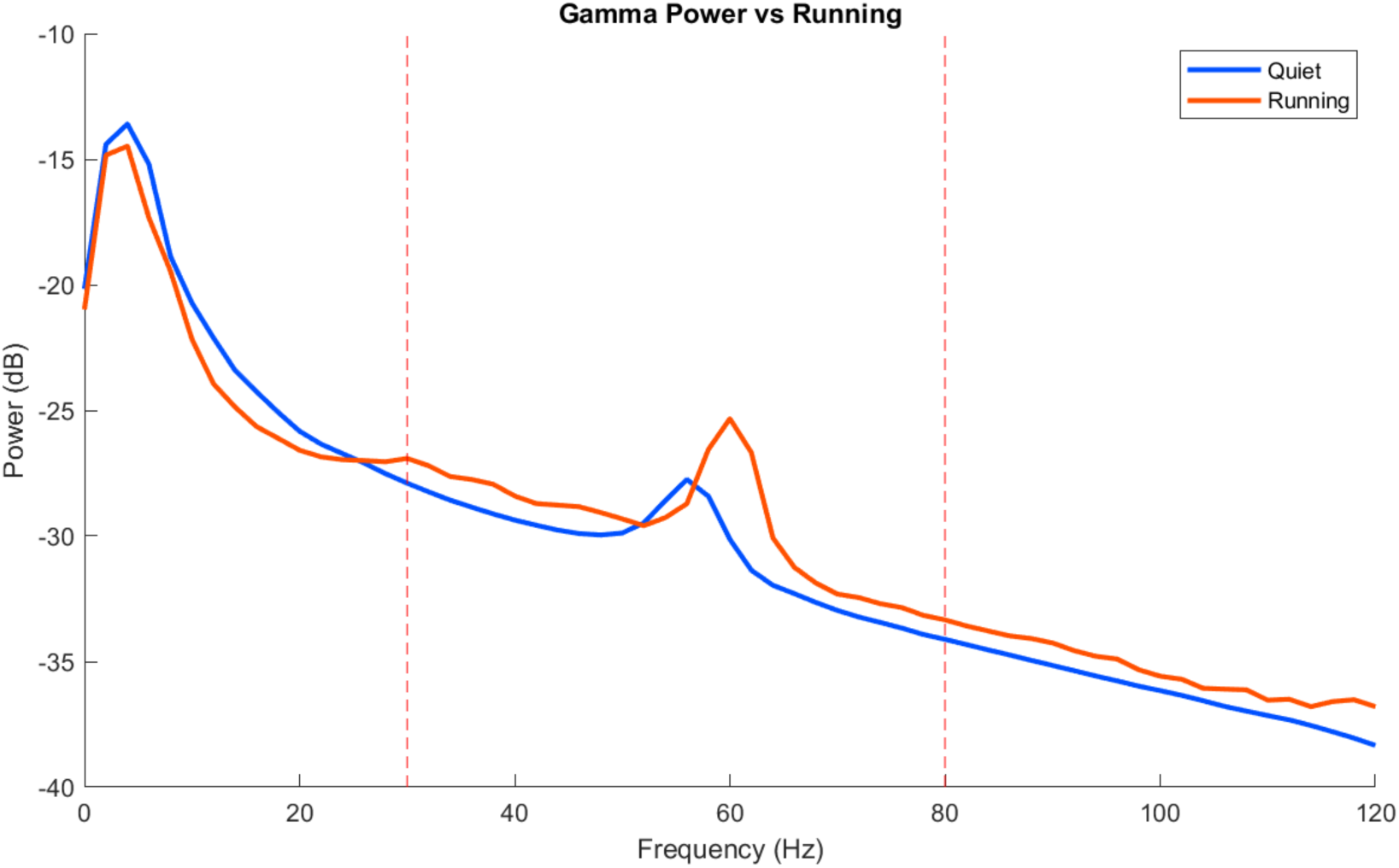

This is often interpreted as an increase in sustained oscillatory activity. However, due to the stochastic nature of neuronal dynamics, we reasoned that this might reflect a higher occurrence of discrete bouts of patterned activity (i.e. events) having energy in that frequency band. To test this idea and uncover the network dynamics underlying these events, we developed a method capable of detecting them in the time domain. This method, called ***C****lustering **B**and-limited **A**ctivity by **S**tate and **S**pectro-temporal feature* (**CBASS**), takes advantage of laminarly distributed multichannel LFP recordings to identify spatio-temporal motifs of LFP activity across channels. This identification is based on 2 criteria: **1)** motifs have energy in the frequency band of interest and **2)** their occurrence increases during the selected behavioral state. The method can be divided into 3 steps

1. **Extraction**, where a set of candidate events is obtained from multichannel LFP recordings in the frequency band of interest
2. **Probability scoring**, where we compute a score reflecting the probability of each candidate event to occur during the state of interest based on spectro-temporal features
3. **Thresholding**, where we find a partition between high and low score events that maximize their distance in the spectro-temporal feature space

### Extraction

The first step of CBASS extracts candidate network events in the selected frequency band and represents them in a parametric space. Each channel of the LFP (Fig. 2 left) is band-pass filtered in the frequency band of interest (Fig. 2 center). In our case, we used zero-phase digital filtering with a 2nd order Butterworth filter (Matlab functions *filtfilt* and butter or their corresponding functions in the Scipy package). Then, we compute the analytical representation of the filtered signal (Matlab function *hilbert* or the corresponding function in Scipy). The analytic representation of a real signal **s(t)** is a complex sequence **s_a(t)** given by:

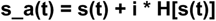

where **H[s(t)]** is the Hilbert transform of **s(t)**. Thus, the real part of the analytical signal is the signal itself and its imaginary part is given by its Hilbert transform. For a band limited time series like the filtered LFP, **s_a(t)** has the very useful properties that its norm and complex argument respectively correspond to the instantaneous amplitude envelope and instantaneous phase of **s(t)** (the norm can be computed with the Matlab function *abs* and the complex argument with the function *angle*. Corresponding functions can be found in the Numpy package). Thus, the analytical signal gives a rich representation of LFP activity at the band of interest and eliminates frequency redundancy problems related to the Fourier transform [1].

**Figure 2.**
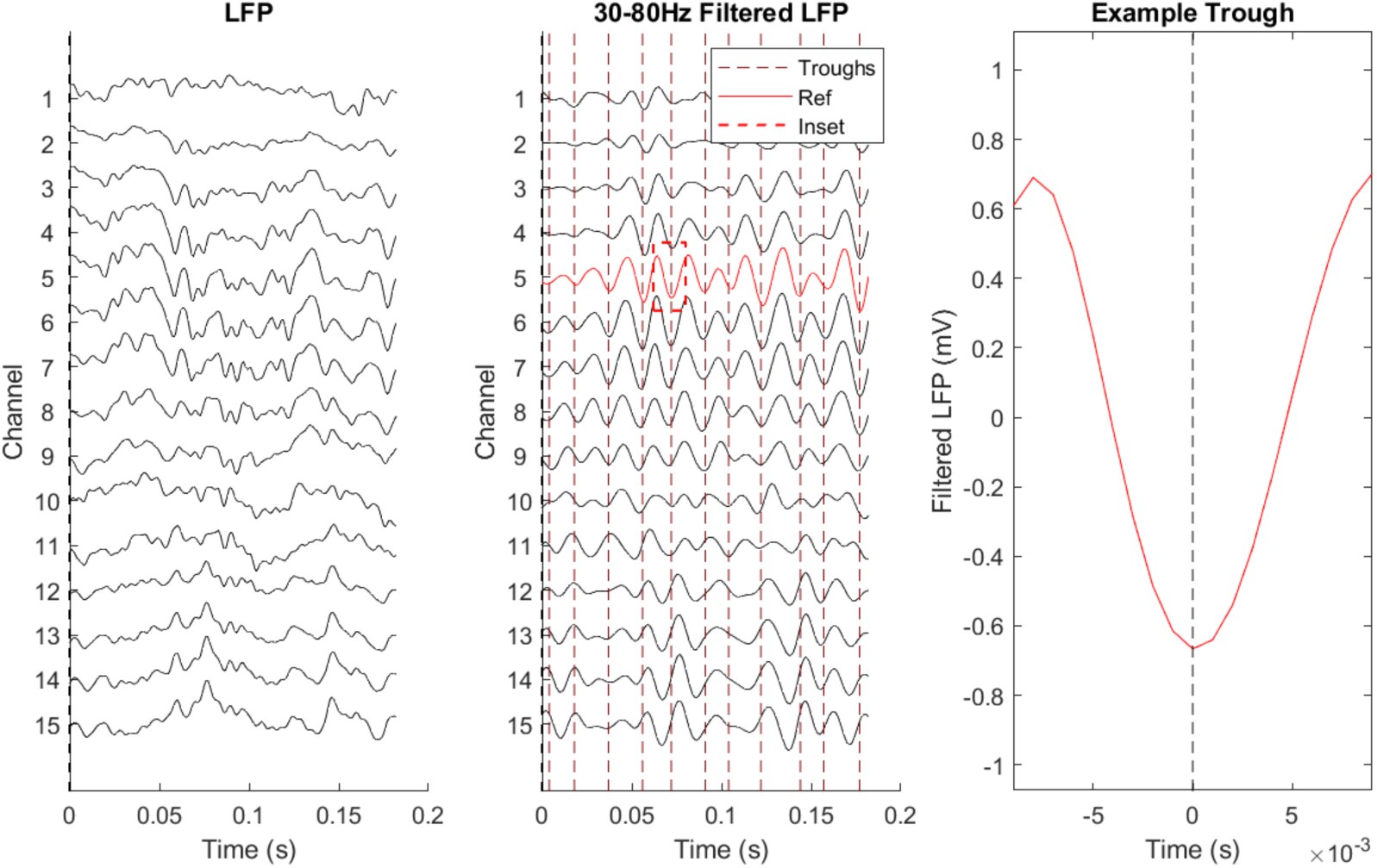

To constrain this representation and make it more amenable to clustering, we select the time points (i.e events) corresponding to the trough of band-passed activity in a reference channel (Fig. 2 center). Troughs are the time points where the argument of the analytical signal (i.e., the phase) is π-valued. Each event is then represented in a parametric space where parameters correspond to the real and imaginary parts of the analytic signal in each channel. Thus, the position of each event in this parametric space gives information about the amplitude and phase of LFP in each channel at the time of troughs in the reference. This offers a comprehensive but constrained representation of the propagation of LFP activity across channels in the band of interest. In our case, the reference was chosen as the channel closest to 400µm of cortical depth (i.e. layer IV, Fig. 2 center - red channel). Different choices of reference did not affect the qualitative outcome of the procedure but resulted in motifs being shifted in time reflecting the propagation of activity between channels. To follow usual conventions in clustering, we designate the data matrix containing the position of each event in the parametric space as **X**. Each element **X(i, j)** corresponds to the value of parameter **j** for event **i** (Fig. 3).

**Figure 3.**
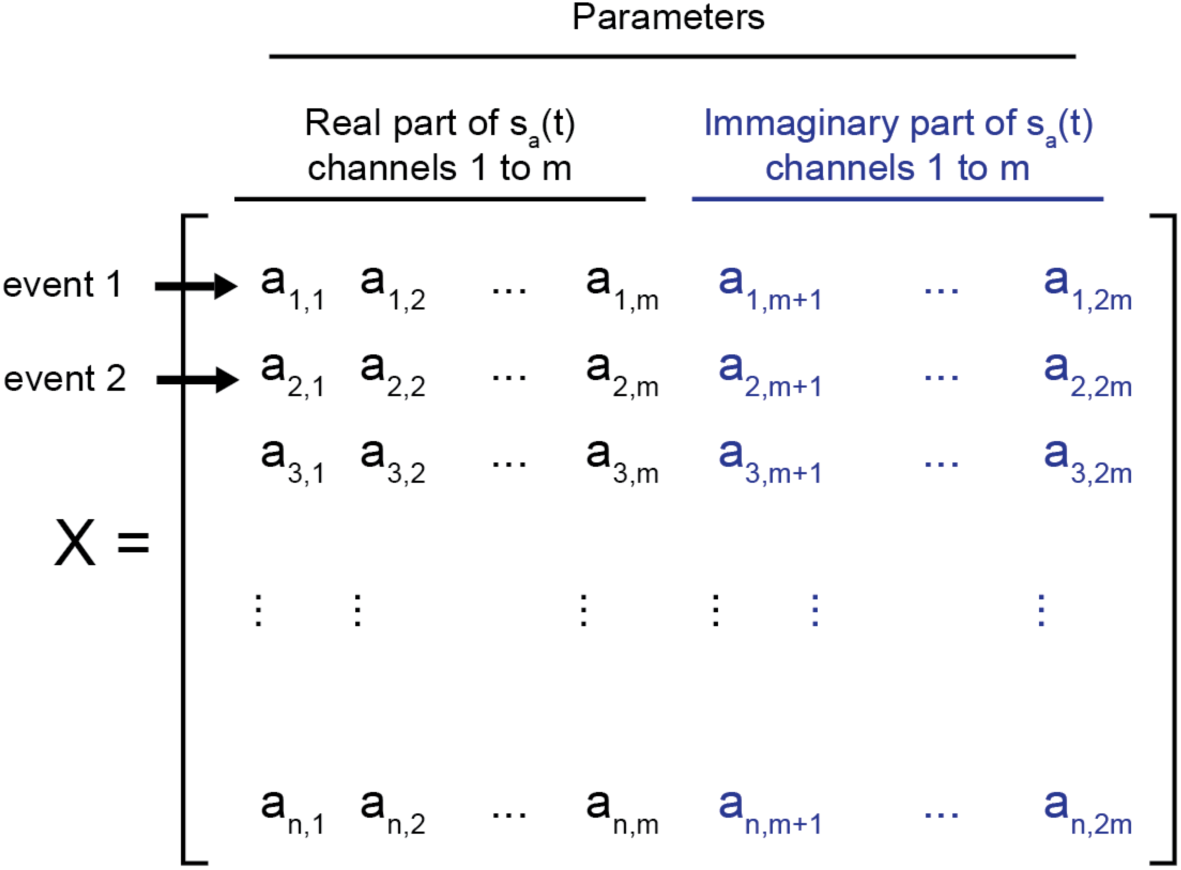

### Probability scoring

We then seek to estimate how likely it is for an event to fall in a region of **X** where the state of interest happens more than by chance. **X** can be conceived as a manifold in a parametric space (Fig. 4 left). Our goal is thus to map variations in the probability of occurrence of the selected state over this manifold. To achieve this, we repeat the following steps

1. The manifold is first partitioned into an arbitrary number k of clusters using the initialization step of the k-means algorithm. Briefly, k centers are drawn at random from the events in **X** All events are then grouped according to which center lies closest to them. These clusters can be thought of as non-overlapping regions of the feature space.
2. We then compute the rate **r** of events occurring during the state of interest in each cluster and compared to **r_all** (i.e. the rate over all events in **X**) using a binomial test of order one. Clusters are considered significantly enriched if **r** is above **r_all** and the binomial test’s p-value is under 0.0001.

After repeating these steps a sufficient number of time (typically 1000 or higher), we compute the enrichment score **s(i)** as the fraction of iterations element **i** was assigned to a cluster where state occurrence was higher than chance. This produces a smooth distribution of score values over the feature space (Fig. 4 center). The number of clusters used has a small but noticeable impact on the result of this procedure. Lower cluster number will produce more smoothing. Conversely, higher cluster numbers will produce distributions having higher entropy at the expense of slower computation time. In our hands, any number between 5 and 100 clusters is acceptable and all give comparable results (Fig. 7; see section potential caveats below for discussion). When needed for visualization or illustration of the different step of the procedure, projections of the **X** manifold to a low dimension space (2D or 3D) are obtained via dimensionality reduction with UMAP[2,3] or PHATE[4].

**Figure 4.**
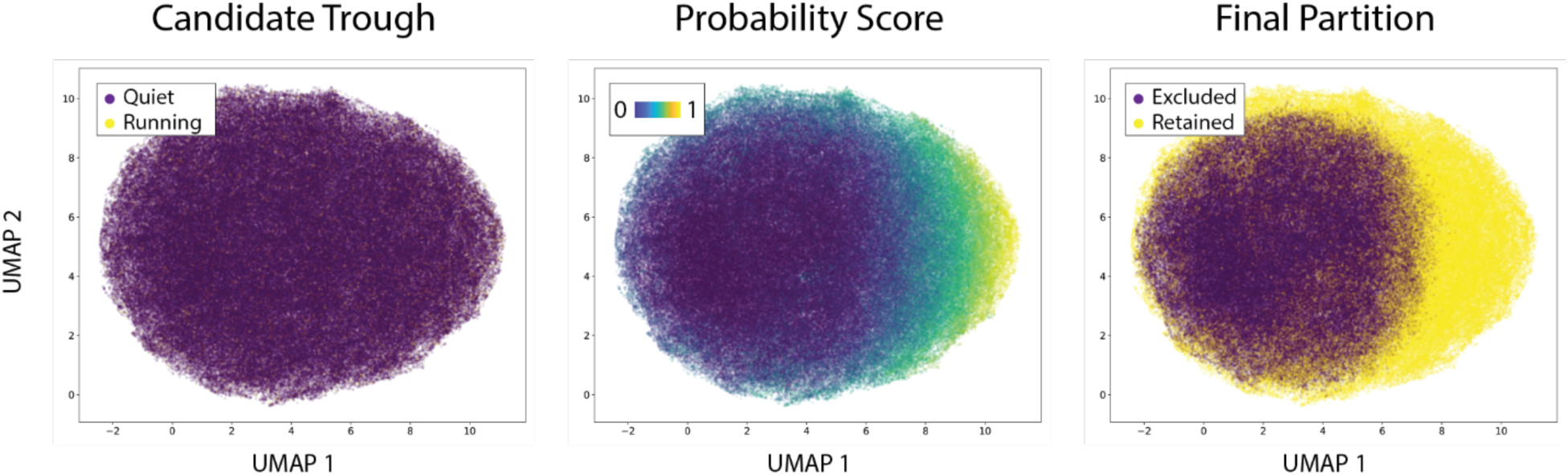

### Thresholding

The last step of CBASS seeks to partition a group of events having homogeneous spectro-temporal features and a high probability of occurring during the state of interest (Fig. 4 right). To achieve this, we find the threshold value of **s(i)** that maximizes the following quantity:

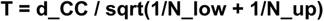

where **d_CC** is the Mahalanobis distance between centroids above and under threshold and **N_low** and **N_up** are the number of events under and above threshold. **T** can be thought of as an analog of the student *t* statistics in multidimensional spaces. Here, searches of the value of **s(i)** maximizing **T** are implemented using the simplex method (Matlab function *fminsearch* or the Scipy function *fmin*).

## Appendix - Generation of surrogate data

To estimate chance level for event detection, CBASS generates surrogate data having the same covariance matrix and the same spectral density in each channel as the original signal (Fig. 5). The LFP is first decomposed into principal components (Matlab function *pca* or corresponding function in the Sklearn package). We then compute the Fourier transform of each principal component (Matlab function *fft* or corresponding in Scipy). The phase of the transform of each principal component is then randomized and a real signal is reconstituted using the inverse Fourier transform (Matlab function *ifft* or corresponding in Scipy). Finally phase randomized principal components are remixed using the principal components loading. This procedure preserves LFP statistics while randomizing spatio-temporal patterns of propagation across channels (Fig. 5).

**Figure 5.**
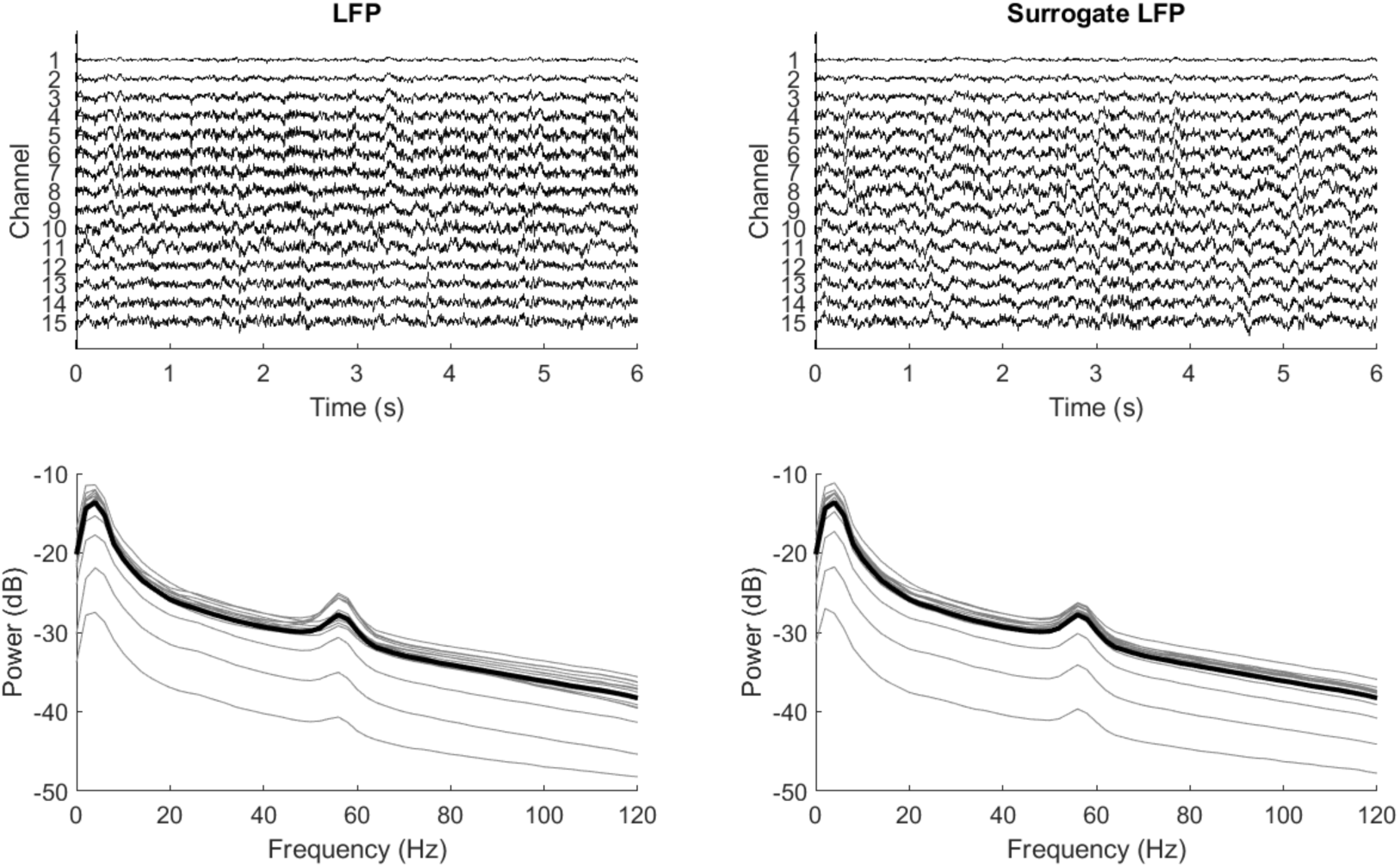

### Validation

The significance of CBASS’s output can be evaluated by comparison to its output on surrogate data (see Generation of surrogate data). We implemented two statistical tests. First, the distribution of enrichment scores between real and surrogate data is compared using the Kolmogorov-Smirnov (KS) test (Matlab function *kstest2* or the Scipy function *ks_2samp*). Failure to pass this test indicates that spectro-temporal features do not yield more information about the occurrence of the state than expected by chance. Second, we calculate the proportion of event in surrogate data falling over the enrichment score threshold. This can be seen as a p-value representing how likely it is for events to be detected when spectro-temporal features do not give information about state occurrence. In the visual cortex of awake mice, we found CBASS to be effective at detecting band specific activity motifs evoked by visual stimulation in the beta range (15-30Hz) and by locomotion in the gamma range (30-80Hz). Current Source Density analysis (CSD) revealed that state enriched events are associated to specific current sink patterns across cortical layers (Fig. 6 left). The frequency of occurrence of the motifs increases during the selected state (Fig 6. center). Finally, spectra acquired when the frequency of occurrence of the motif is high (Fig. 6 right) look very similar to spectra evoked by the selected state for each type of activity (Fig. 1).

**Figure 6.**
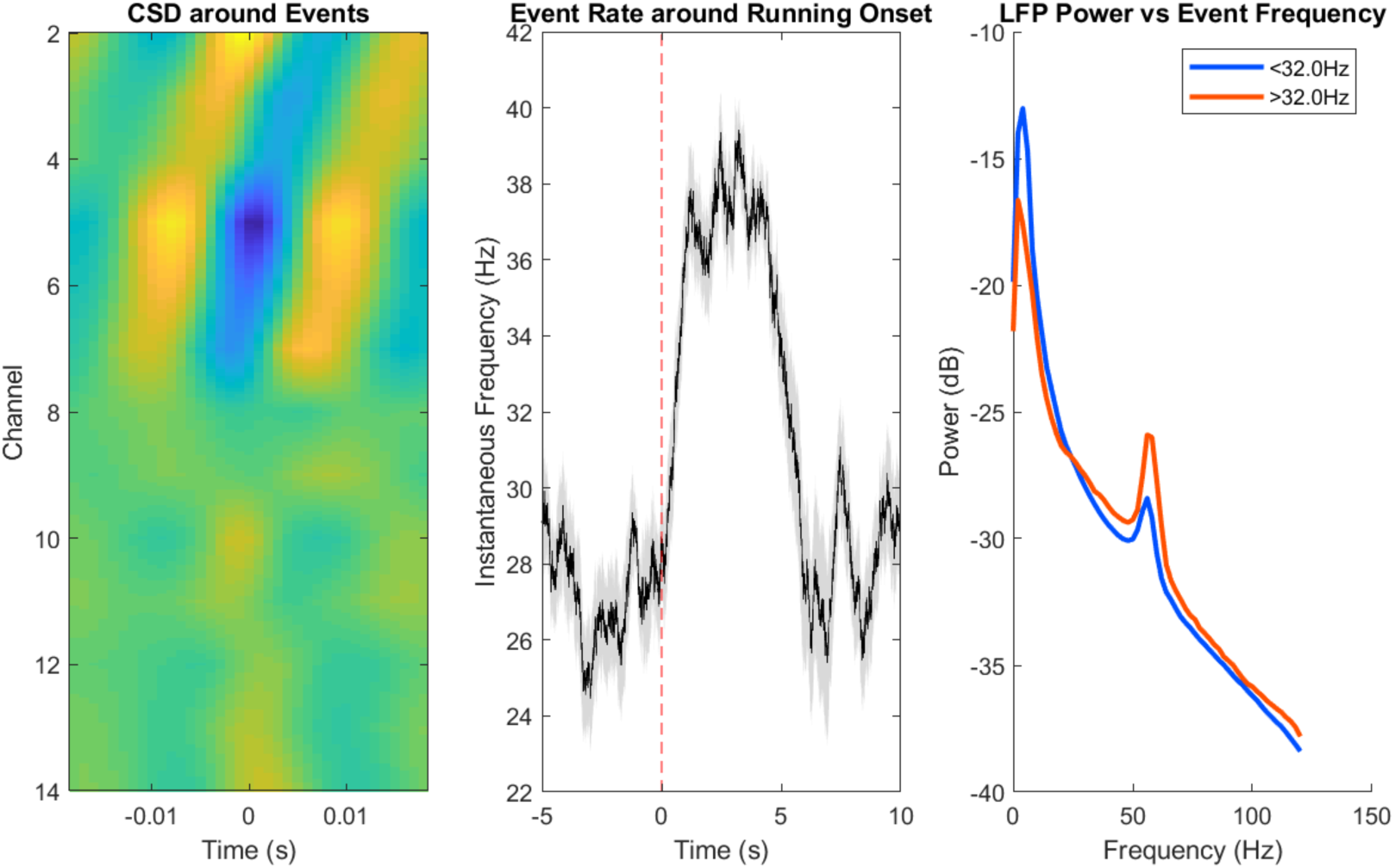

### Potential caveats - optimizing cluster number for probability estimation

To estimate state occurrence probability based on features, CBASS partitions a set of band specific events into an arbitrary number of clusters (see section Probability scoring above). This segmentation is repeated to produce a smooth probability distribution of the state of occurrence over events based on spectro-temporal dynamics. Lower cluster numbers will result in more smoothing whereas higher number will tend to produce more contrasted distributions. If the number of clusters is not sufficiently high, the procedure might fail to detect small regions where state probability is high. Choosing a number that is too high will increase computation times. In our hands, the output of the method is very robust to changes in cluster number (Fig. 7). However, applying CBASS might require testing an increasing number of clusters for the kind of problem that is meant to be addressed. We advise choosing the minimal number of clusters that yield a stable result.

**Figure 7.**
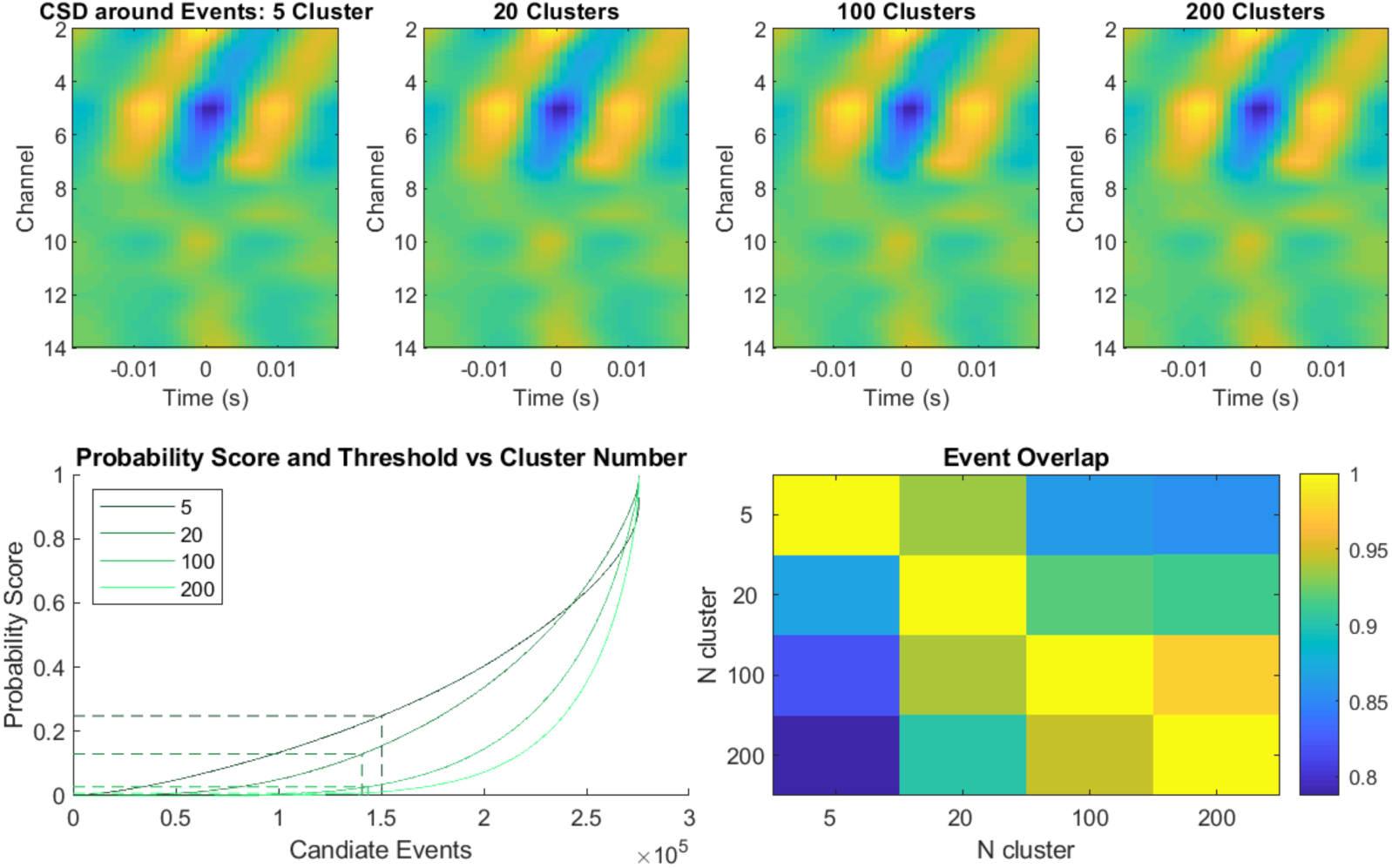

---

**Supplemental Table 1.**
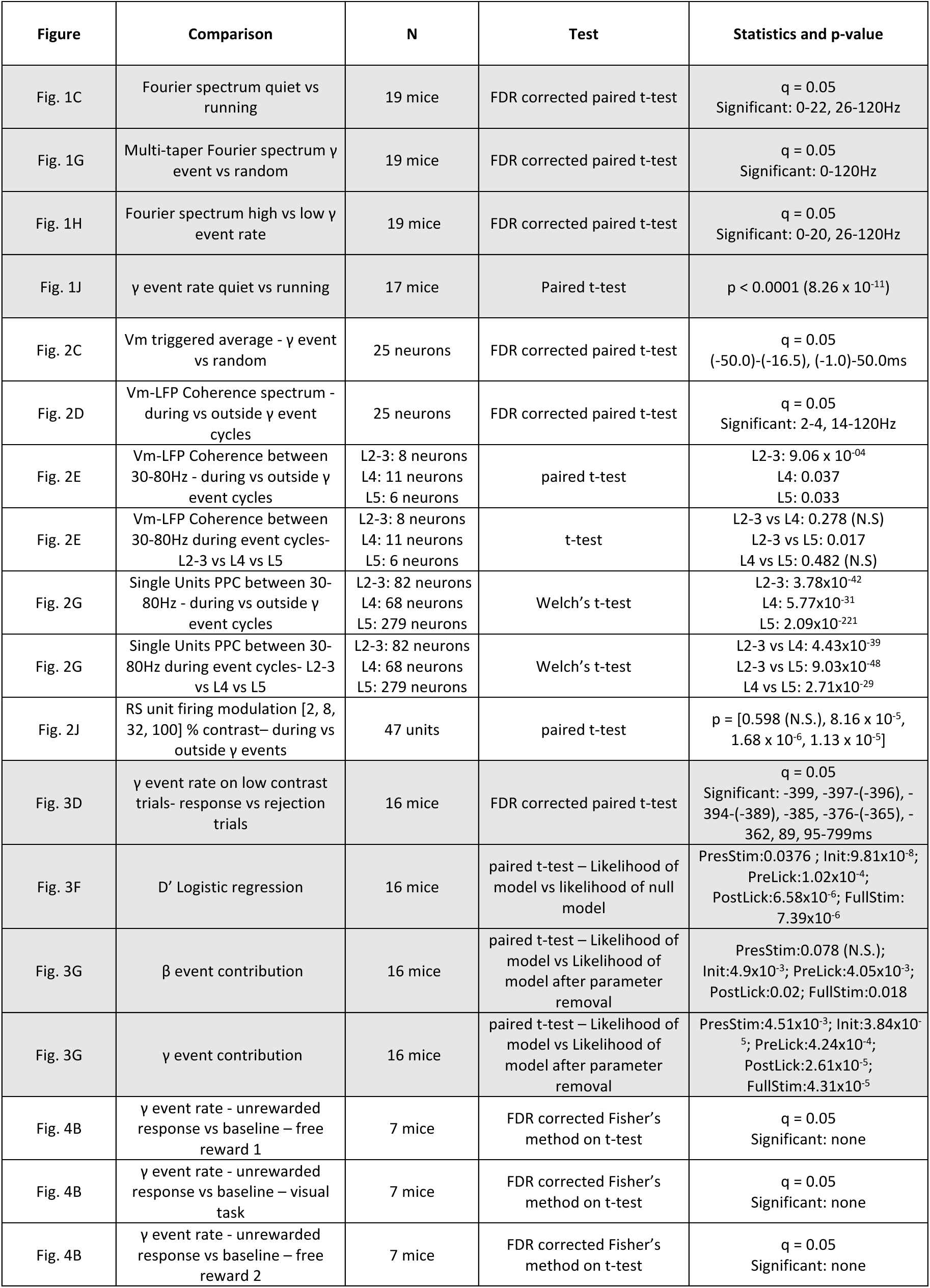
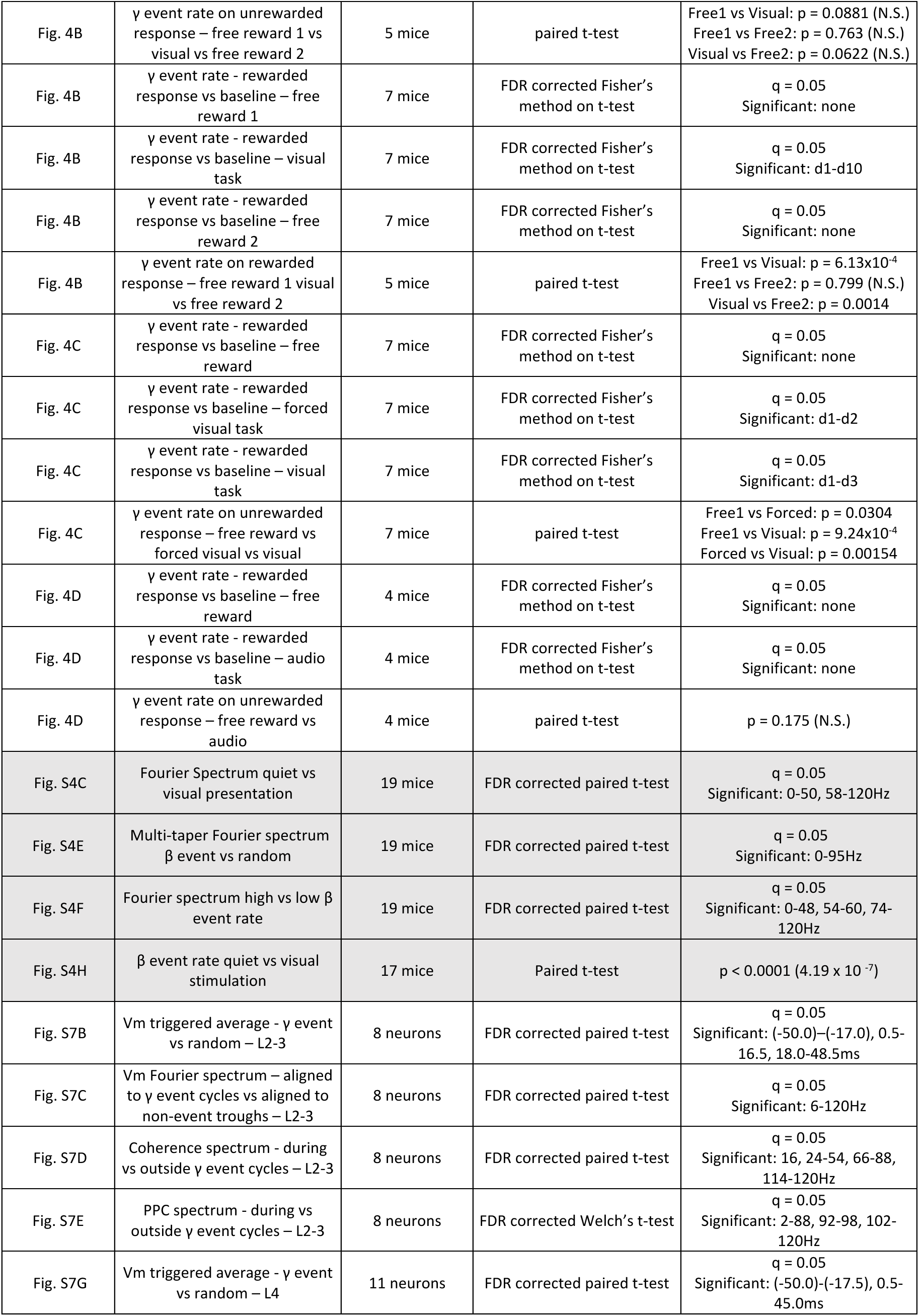
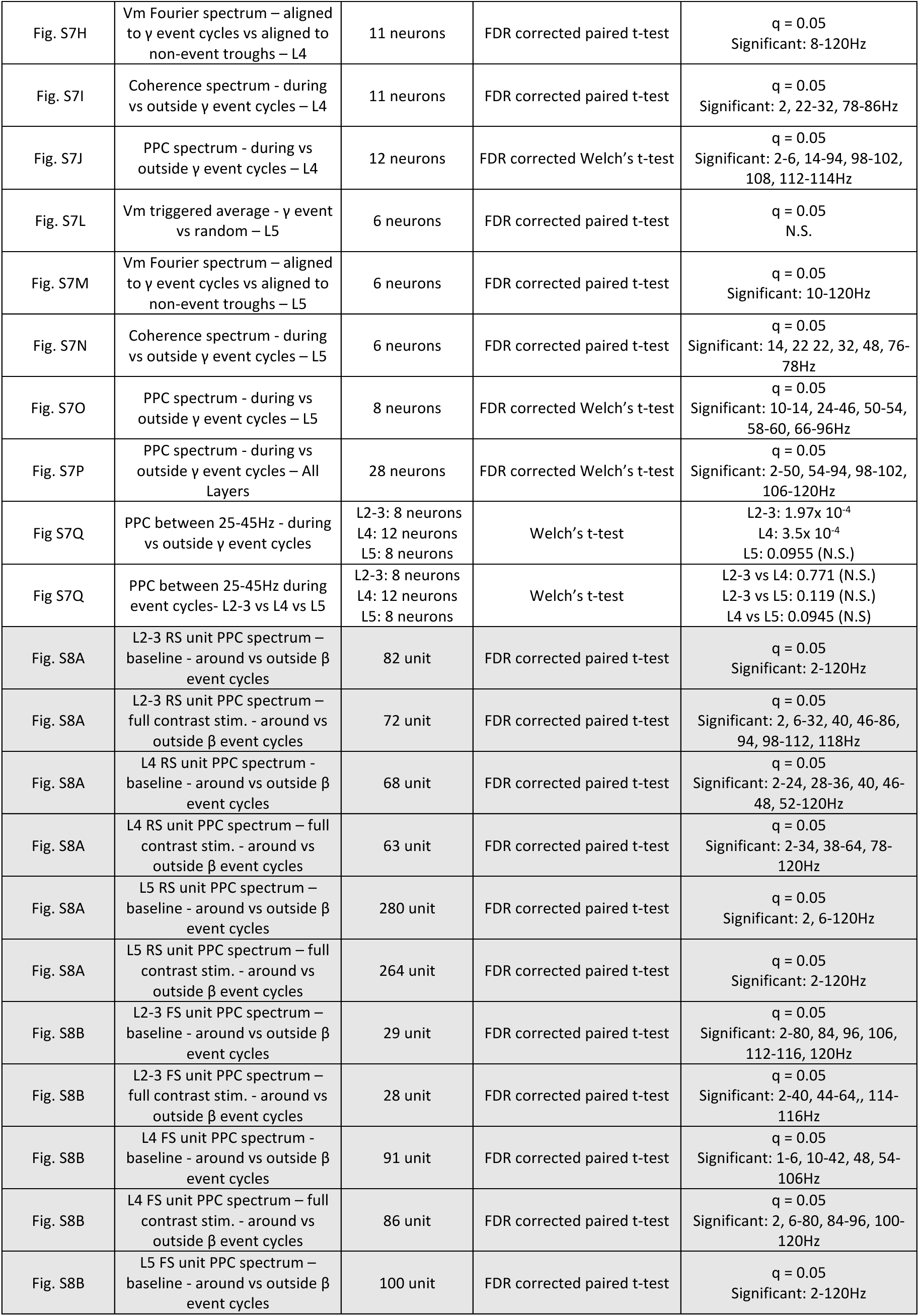
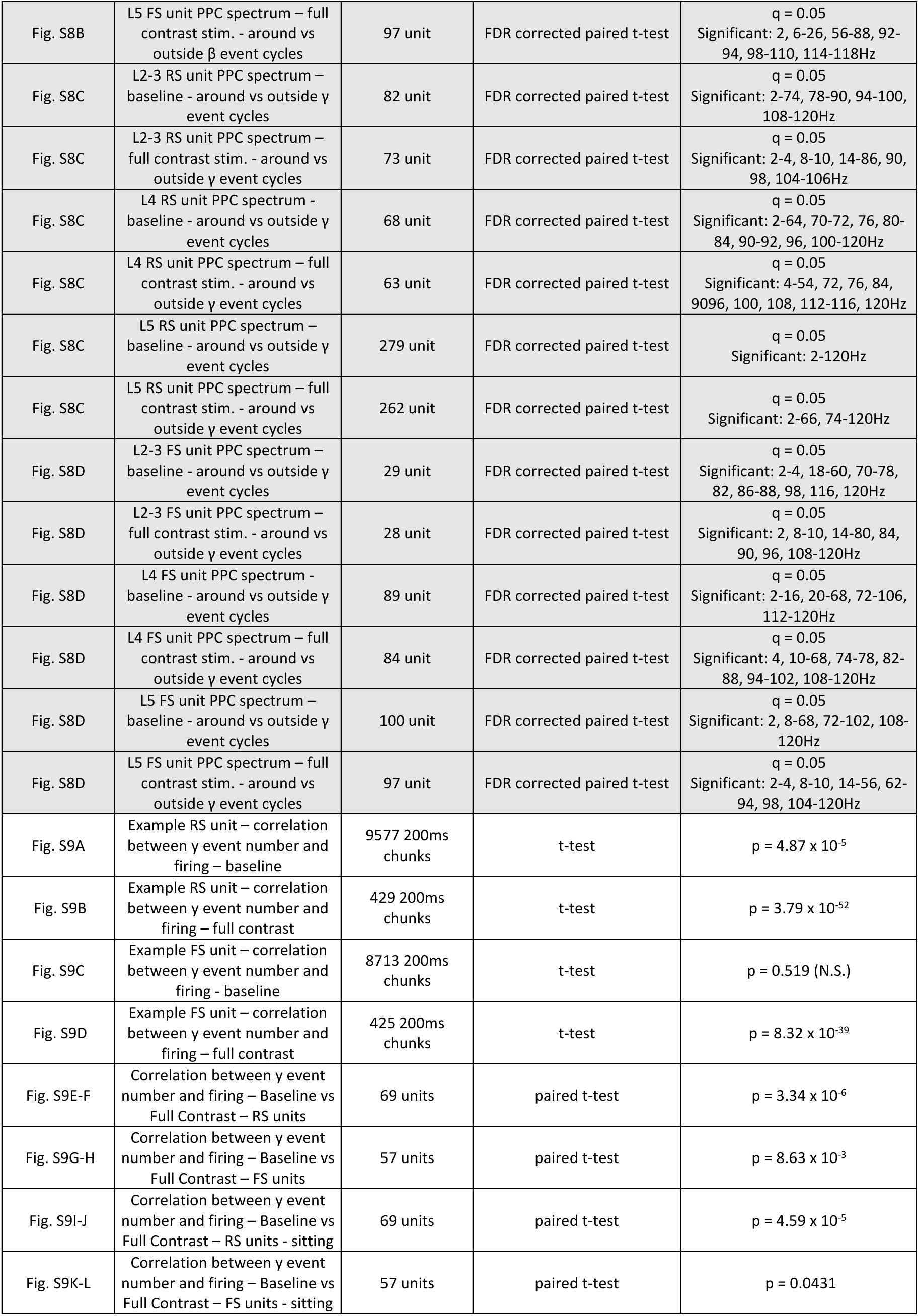
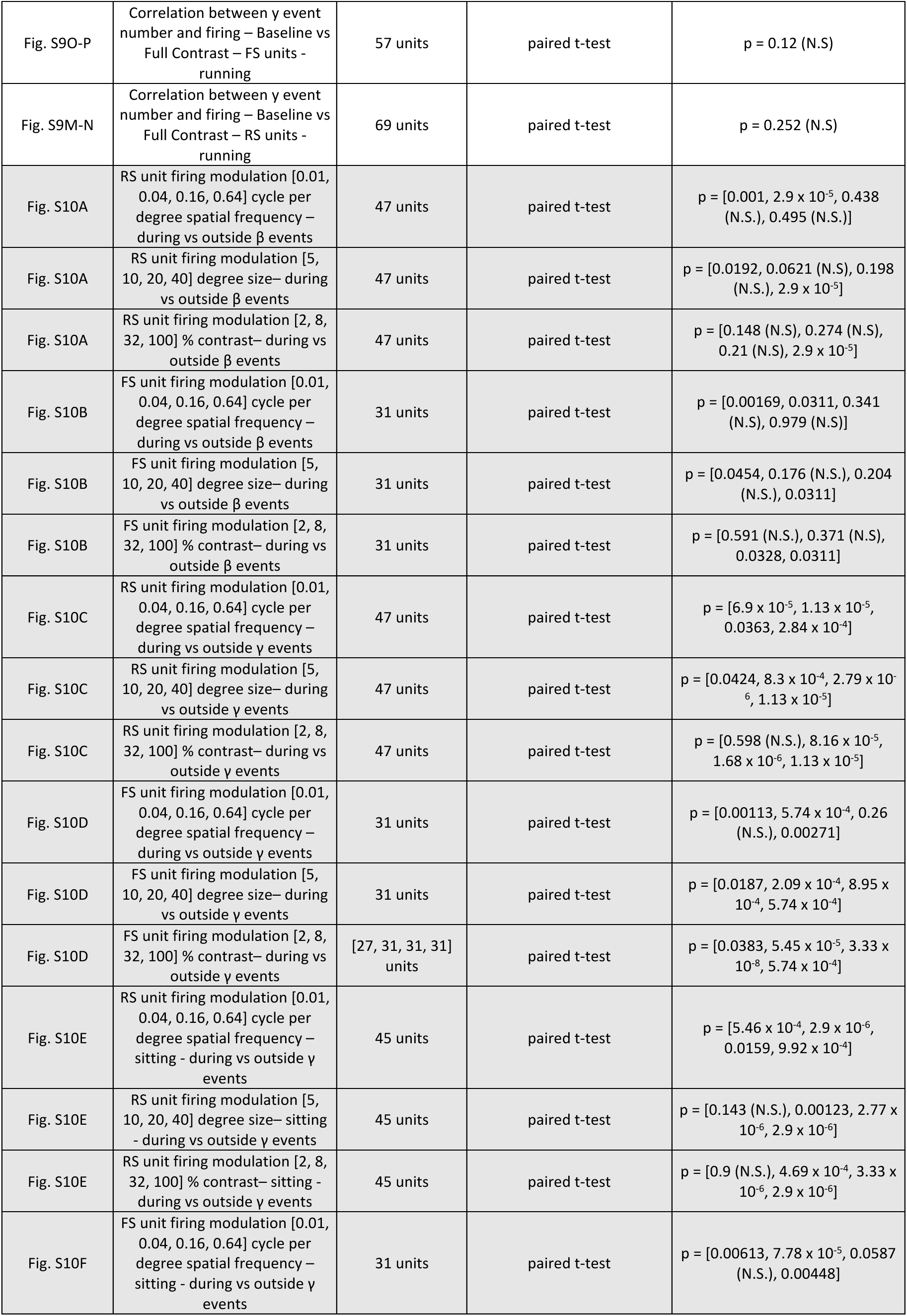
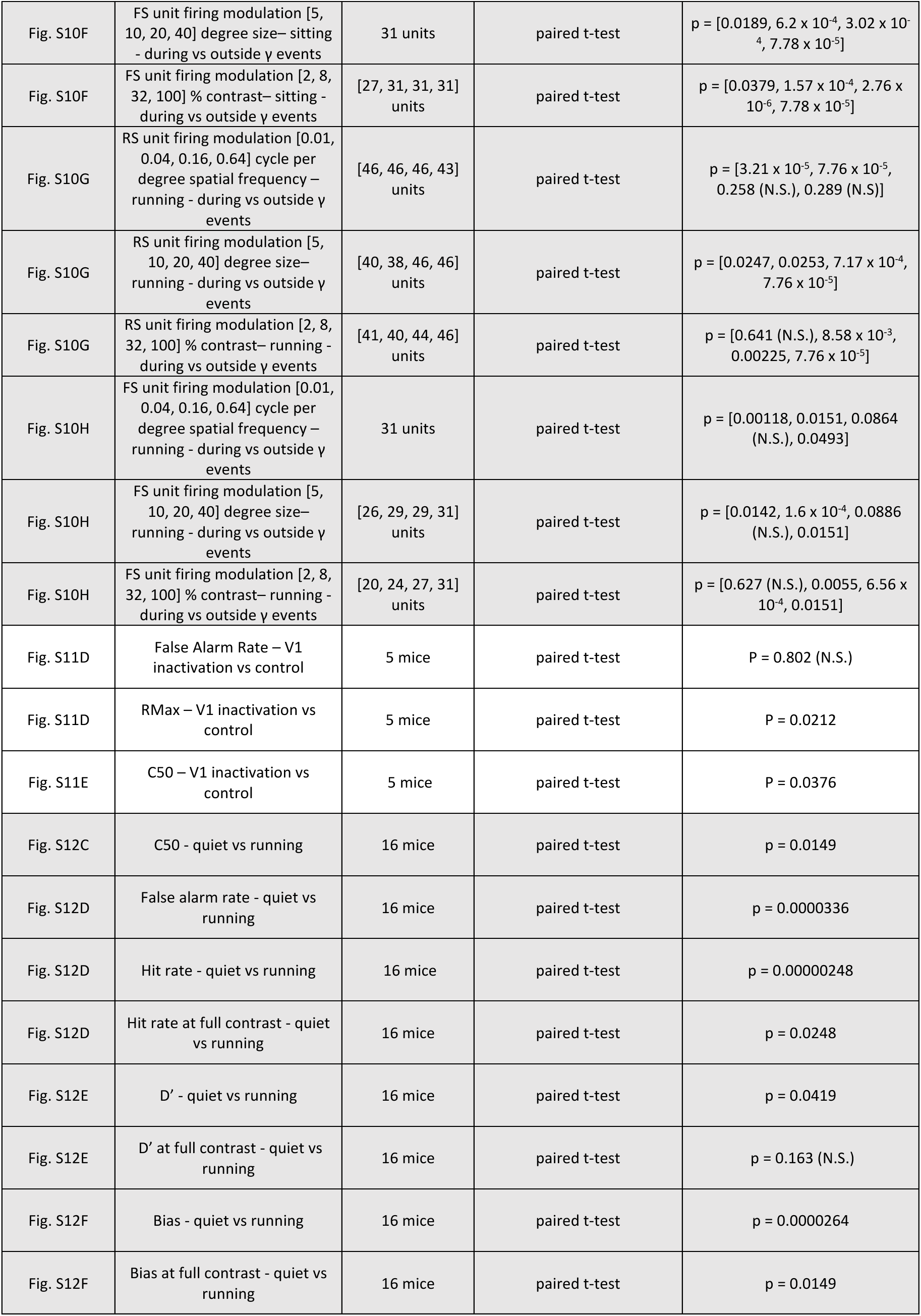
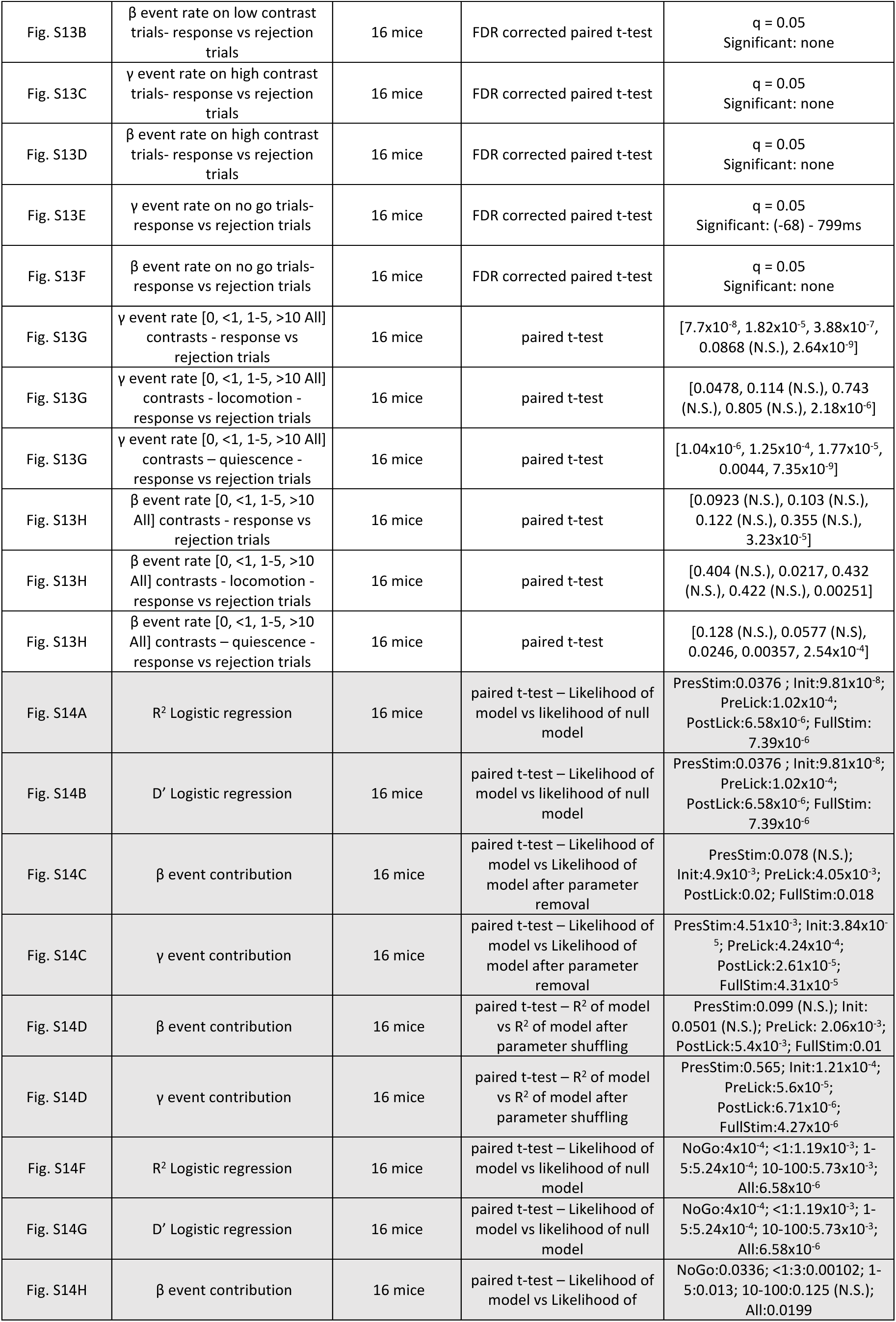
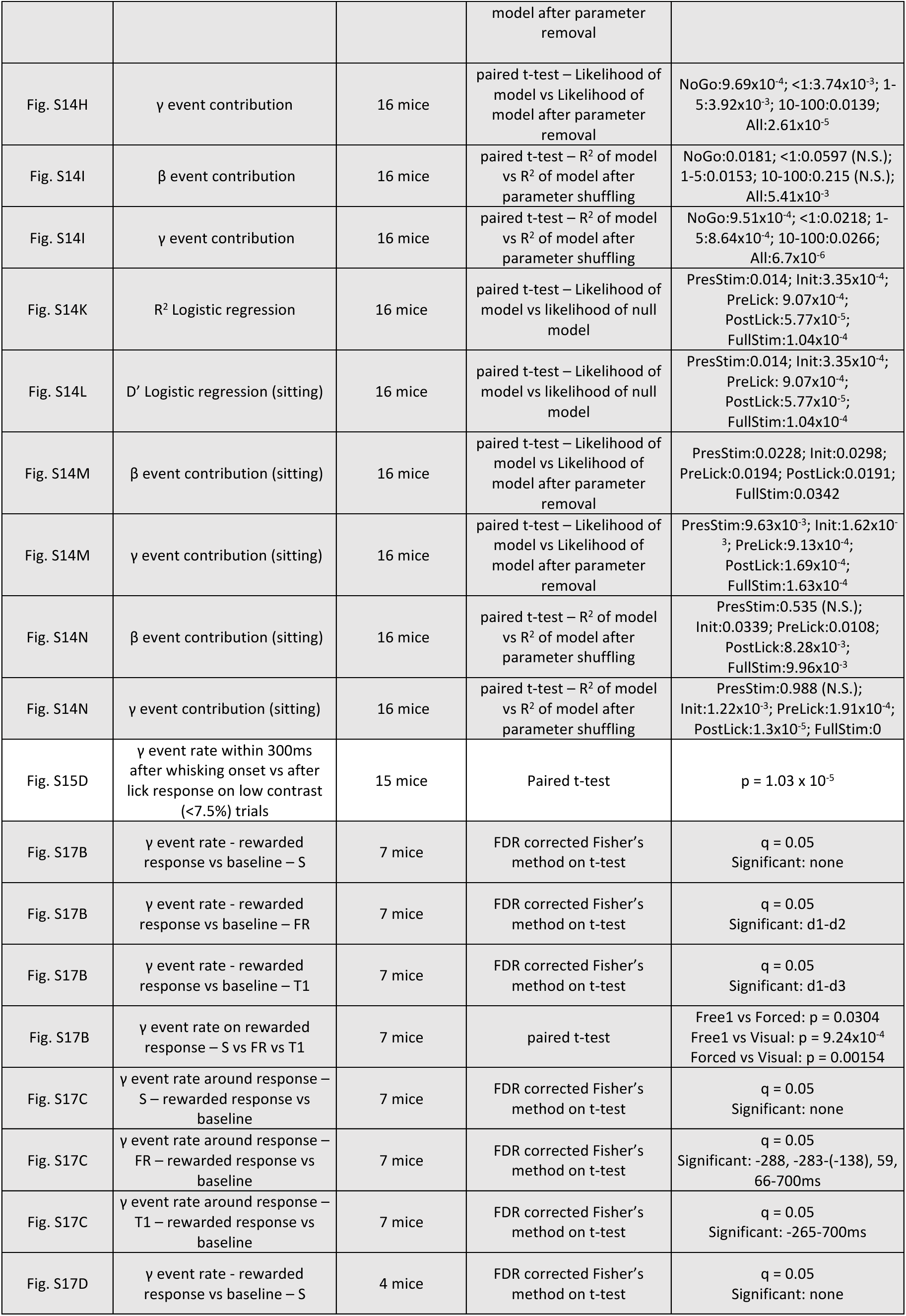
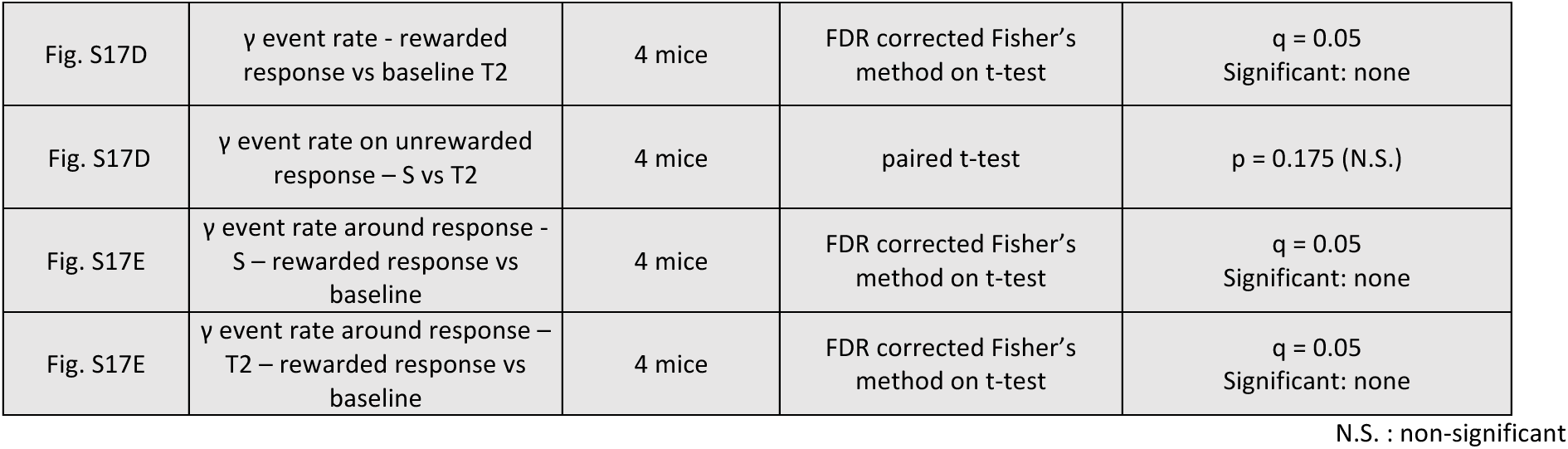
Summary of all statistical analyses.

